# Finding and extending ancient simple sequence repeat-derived regions in the human genome

**DOI:** 10.1101/697813

**Authors:** Jonathan A. Shortt, Robert P. Ruggiero, Corey Cox, Aaron C. Wacholder, David D. Pollock

## Abstract

**Background:** Previously, 3% of the human genome has been annotated as simple sequence repeats (SSRs), similar to the proportion annotated as protein coding. The origin of much of the genome is not well annotated, however, and some of the unidentified regions are likely to be ancient SSR-derived regions not identified by current methods. The identification of these regions is complicated because SSRs appear to evolve through complex cycles of expansion and contraction, often interrupted by mutations that alter both the repeated motif and mutation rate. We applied an empirical, kmer-based, approach to identify genome regions that are likely derived from SSRs.

**Results:** The sequences flanking annotated SSRs are enriched for similar sequences and for SSRs with similar motifs, suggesting that the evolutionary remains of SSR activity abound in regions near obvious SSRs. Using our previously described P-clouds approach, we identified ‘SSR-clouds’, groups of similar kmers (or ‘oligos’) that are enriched near a training set of unbroken SSR loci, and then used the SSR-clouds to detect likely SSR-derived regions throughout the genome.

**Conclusions:** Our analysis indicates that the amount of likely SSR-derived sequence in the human genome is 6.77%, over twice as much as previous estimates, including millions of newly identified ancient SSR-derived loci. SSR-clouds identified poly-A sequences adjacent to transposable element termini in over 74% of the oldest class of *Alu* (roughly, *AluJ*), validating the sensitivity of the approach. Poly-A’s annotated by SSR-clouds also had a length distribution that was more consistent with their poly-A origins, with mean about 35 bp even in older *Alus*. This work demonstrate that the high sensitivity provided by SSR-Clouds improves the detection of SSR-derived regions and will enable deeper analysis of how decaying repeats contribute to genome structure.

## Background

Simple sequence repeats (SSRs) are 1-6 bp tandem repeats that have been estimated to comprise 3% of the human genome (1, 2). SSRs are notable for their unusual mutation process; after they reach a threshold length (3-5 tandem motif repeats), the rate of slippage during DNA replication dramatically increases, resulting in rapid expansion or contraction of SSR loci. These events may occur at a rate of 1 x 10^-3^ per locus per generation (3, 4), many orders of magnitude faster than point mutation rates, and can modify structural and regulatory functions, contributing to disease (5). In addition, because they are enriched in promoters, highly mutable, and provide a rich source of heritable variation, SSRs were proposed to be evolutionary “tuning knobs” (6–10). Numerous recent studies have highlighted the potential functional role of SSRs in gene regulation (11–14) and a better understanding of SSR evolution may therefore allow insights into how function can arise from constantly changing genomic structure.

A proposed life cycle for SSRs includes intertwined stages of birth, adulthood, and death (15–18). *De novo* birth of an SSR at a location occurs when a short series of repeats arises by chance mutations, and aided and extended by the tendency of duplications to occur via normal (non-SSR) slippage events that result in tandem duplication of short motifs (15, 18). If the number of simple sequence repeats exceeds some threshold length, which can depend on the composition and purity of the repeated motif (19), then the probability of slippage will increase with a slight bias towards increasing numbers of repeats (4, 20–22). Additionally, although there is a clear lower bound on repeat lengths (zero, obviously) and the slippage rates for small numbers of repeats is low, there is no upper bound on repeat lengths unless it is biologically imposed. These factors together are thought to result in rapid expansion in the number of motifs at SSR loci and suggests that accurately describing the length and distribution of SSRs may provide a new source of insights into genome biology.

It is thought that during SSR “adulthood”, slippage-induced expansions and contractions (usually one repeat at a time) can rapidly alter the length of SSR loci, but mutations that disrupt the composition of tandem repeats also accumulate and slow or stop the slippage process (23, 24). The SSR life cycle is potentially complicated by rare multiple-motif copy number mutations that are thought to be biased towards large deletions, and by selection against long repeat lengths that may lead to upper size limits (20, 21, 25). Transposable elements (TEs) also contribute to SSR generation by introducing pre-existing repeats at the time of TE replication, by introducing poly-A tails (in the case of retroelements), or by repeatedly introducing sequences that are likely to give birth to new SSRs (16, 26, 27).

SSR death presumably occurs after either sufficiently large deletions at a locus have occurred or after enough mutations have accumulated so that there are no longer uninterrupted tandem motif stretches above the threshold length (17). After the death of an SSR, remnants of the formerly active SSR locus may remain in the genome, sometimes spawning an active SSR locus (with the same or similar motif) capable of expansion by slippage; this phenomenon has been observed but not characterized in great depth (15).

The abundance of active SSRs in the genome and their finite lifetime suggest that dead SSRs may also be abundant, although their high slippage mutation rate and complex, motif-dependent evolution makes modeling their evolutionary outcomes difficult. The identification of dead SSRs remains important if for no other reason than because their presence in the genome can confound the detection and annotation of other genomic elements (28). Several reports have noted that the sequence composition near SSRs is biased towards the adjacent SSR motif, and it has been proposed that such sequences are SSR-derived (29, 30); however, the origin of this biased sequence has not been explored in detail. Part of the problem is that Tandem Repeats Finder (TRF) (31), the current predominant method for finding genomic repeats, although mathematically elegant and computationally efficient, is designed to detect perfect and near-perfect repeats, and provides little information about more degenerate SSR-derived loci. The ability to better identify degraded SSRs at various ages and stages of their life cycle would thus aid in annotation of the genome and inform on the origins and history of regions in the genome where they reside.

Here, we report a new method to detect SSR-derived sequence using a probability-clouds (*P-clouds*) (32, 33) based approach. This approach uses empirical counts of oligonucleotides (oligos) to find clusters (or clouds) of highly enriched and related oligos that, as a group, occur more often than predicted by chance. The *P-clouds* method has been applied to identify various repetitive structures in the human genome (32, 33), including transposable elements, but has not yet been applied to identify SSRs (which were specifically excluded from the original method). The use of empirical oligo enrichment, coupled with alignment-free and library-free detection, makes *P-clouds* both fast and particularly well-suited to annotate regions resulting from the complex mutational processes associated with SSR loci. We obtained sets of p-clouds in regions flanking perfect live SSRs under the hypothesis that such regions will be enriched in the mutated detritus of the SSRs (34). These SSR p-clouds, called SSR-clouds, were then used to re-define the spans of active SSR regions and locate dead SSR loci that were not previously identified. We also provide further evidence that SSRs frequently spawn new SSR loci with similar motifs, presumably because the low sequence degeneracy of SSR detritus regions makes them fertile spawning grounds.

## Results

### Characterization of perfect SSR loci in the human genome

Uninterrupted perfect SSR loci abound in the genome. SSR sequence motifs of 1-6 bp were grouped into motif families comprised of a motif, its reverse complement, and any possible alternate phase of the motif or its reverse complement (e.g., AAC, ACA, CAA, GTT, TGT, and TTG all belong to the same motif family) to create a total of 501 separate SSR motif families. If a longer motif was a repeated multiple of a shorter motif (e.g., ATAT versus AT), that motif was assigned to the shorter motif. The unmasked human genome (hg38) was annotated (Table S2) with these motif families to locate every *perfectly* repeated contiguous SSR locus (one that contains no point mutation, insertion, deletion, or motif phase shift; loci separated by 1 or more bp were assigned different loci in this analysis) at least 12 bp in length. A total of 4,551,080 perfect (uninterrupted) SSR annotations were found, covering 68.8 Mb (∼2.2% of the genome). These perfect repeats constitute over three-quarters (77.8%) of the 88.4 Mb SSR sequence (2.85% of the human genome) annotated using standard TRF settings.

The 12 bp minimum length for SSR loci is consistent with reports that established an SSR expansion threshold cutoff at around 10bp for motifs ≤ 4 bp (15, 38, 39), and is consistent with our own analyses of when perfect SSR frequencies significantly exceed expectations based on genomic dinucleotide frequencies (see Figure S1). The most highly-represented SSR is the mononucleotide repeat poly-A/poly-T (henceforth referred to as just poly-A) with 703,012 separate loci. Consistent with previous reports, many (467,092, or 66.44%) of these poly-A’s overlap with an annotated *Alu* (40), and 536,938 (76.38%) overlap with any annotated transposable element. Some caution is warranted in interpreting this result, both because the poly-A tail and the A-rich region in the center of many *Alus* may or may not contain a perfect repeat, and because RepeatMasker is inconsistent about whether it includes a poly-A tail in a repeat annotation. Nevertheless, this result indicates the minimum extent to which transposable elements contribute to the frequency of poly-A loci in the genome. Other than poly-A, the next most represented motif is CA/TG with 170,729 separate annotations, only 3,206 (1.88%) of which are found in an *Alu* element. Although all possible SSR motifs families have at least one locus in the genome, the most common motif families tend to have much simpler motifs than the least common (64% of the 50 most common motifs contain only 1 or 2 nucleotides, and only three of the most common motifs contain all 4 nucleotides, while 82% of the least common motifs contain all four bases (see Table S2), suggesting more frequent rates of origination for these simpler motifs. There is also an enrichment of shorter motifs amongst the most common SSRs, a trend that is consistent with previous observations (4, 41).

### Characterization of sequence bias in the regions flanking perfect SSRs

Sequence biases in the regions flanking SSRs are a rich resource for understanding the evolutionary remains of SSR activity. Perfect SSR loci are often closer to each other than expected by chance, with an extremely high peak under 10 bp separation, and leveling off before 100bp (Figure S2). Reasonable explanations for close repeats include that they were previously a single locus that was divided by imperfections, or that new repeats were spawned from a single repeat’s detritus. Indeed, the repeated motifs of adjacent SSR loci often share high sequence similarity. The most represented repeated motif near a perfect SSR locus is often the repeated reference motif itself, and other similar motifs are also highly over-represented (Figure 1). As an example of more complex families, we considered (ATGC)_n_ loci, and adjacent SSRs that had 1, 2, or 3 different nucleotides. As with the simpler motifs in Figure 1, similar motifs are highly enriched at short distances from (ATGC)_n_ repeats (Figure 2), while dissimilar motifs are far less enriched. These observations suggest that SSRs can originate from the periphery of existing SSR loci where sequence is already biased towards simple sequences (30). Under this hypothesis, dissimilar families that require multiple mutations to reach a threshold slippage length are found at lower frequencies because they are more difficult to seed.

**Fig. 1.**
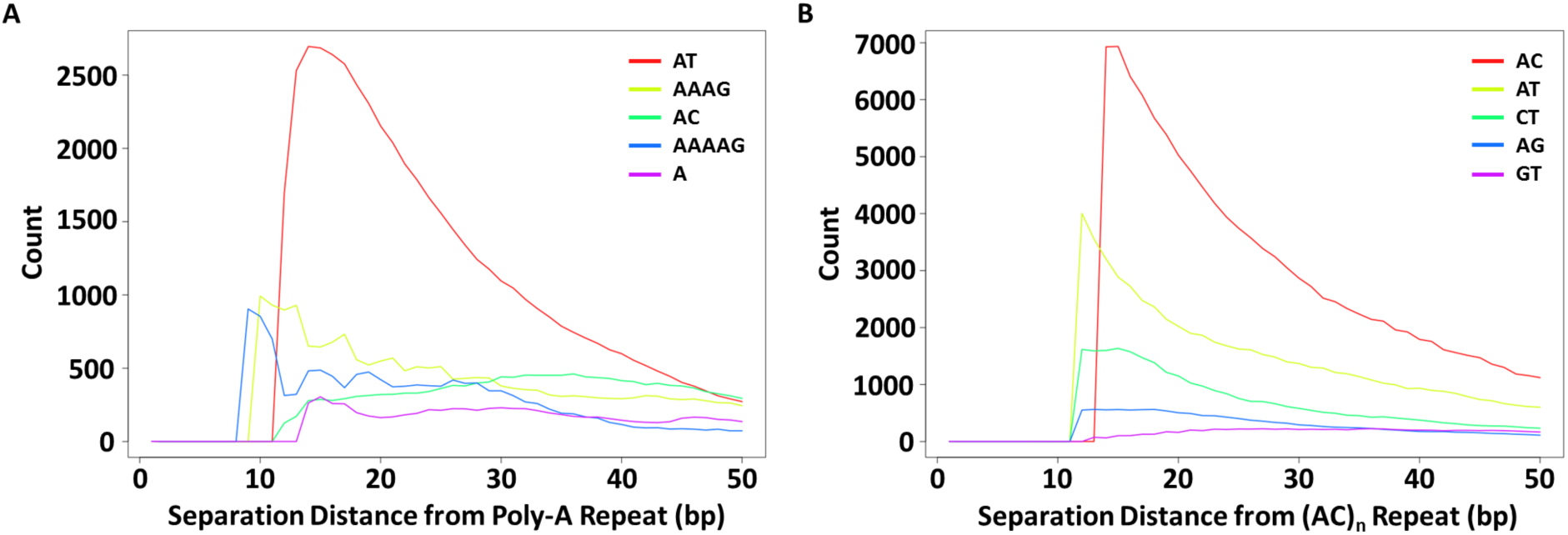
Clustering of SSR loci depending on motif similarity. All perfect SSRs (≥12 bp) were annotated in a transposable-element masked version of the human genome (hg38) and the count of nearby SSR motifs were recorded as a function of distance from the repeat. Here, we show the 5 motifs that are most frequently found near (A) perfect poly-A SSRs (n=350,763); and (B) perfect (AC)_n_ SSRs (n=85,161). The motifs of nearby SSRs often differ from the repeated motif by simple mutations. To allow for overlapping non-reference motif families (i.e., a compound locus comprised of two or more different motif families), x=0 begins 11 bp within the perfect reference motif repeat. Flat curves at x=0 reflects that the first several bases are still part of the perfect repeat and thus can only be annotated by another family to the extent that their motifs overlap.

**Fig. 2.**
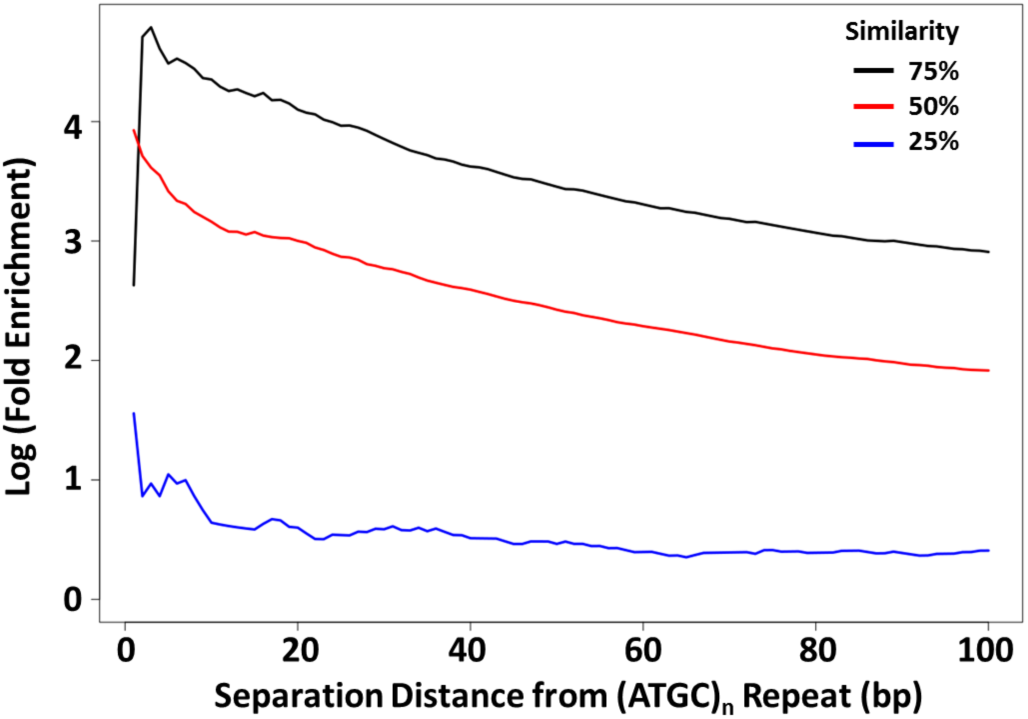
Enrichment of similar SSR loci near ATGC repeat loci. The average enrichment levels of perfect SSR loci within 100 bp of a perfect ATGC repeat locus are shown for SSR families with motifs with 1 (75%, black), 2 (50%, red), or 3 (25%, blue) differences from the ‘ATGC’ motif. Enrichment for SSR motifs was determined relative to the genomic average for all possible motifs with the given difference.

To better describe the extent of the periphery around SSRs, which is known to deviate from random sequence (29, 30) and may represent a detritus field of mutated repeats (34), we measured similarity to each repeated perfect motif within 200 bp on either side of the repeat. There are differences depending on the size and repeat motif, but in general similarity extends at least 50-100 bp on either side of motifs (Figure 3). This size of detritus field is consistent with the idea that regular SSR seeding occurs from this detritus. As a side note, poly-A sequences had detritus fields on their 3’ side, but not their 5’ side, because they commonly originate from transposable elements (Figure S3) whose uniform sequence obscured the presence of detritus fields.

**Fig. 3.**
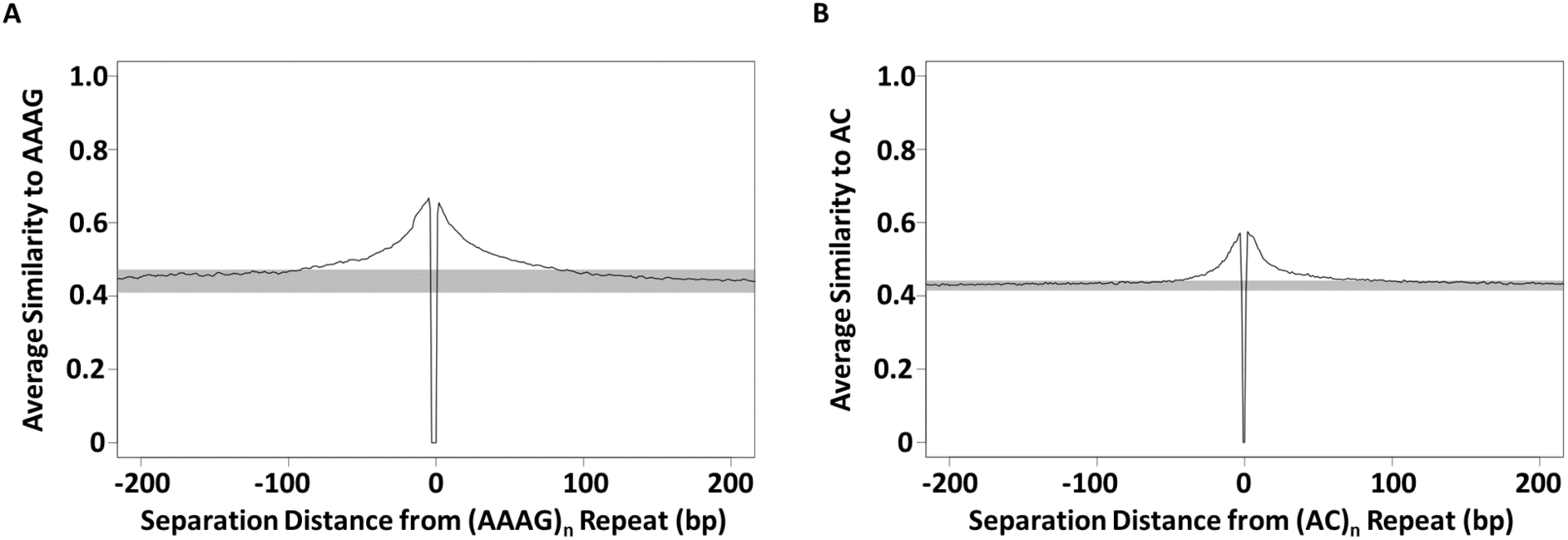
Decay of sequence similarity with distance from perfect SSR repeats. Average similarities were calculated for short segments within 200 bp of perfect SSR repeats with a given motif. Similarity was measured as the proportion of identical nucleotides at each position for a segment of the same length and read direction as the repeated motif shown, (AAAG)_n_ in A, (AC)_n_ in B. For example, a segment reading “ATAG” would have a similarity of 0.75 with the repeat motif “AAAG”. Average similarities were calculated for segments beginning at every nucleotide separation distance within 200 bp of the perfect repeat beginning or end. The black line shows the average similarity to each repeat, while the gray box shows a range of 3 standard deviations from the mean similarities calculated in 700 bp windows from 300-1,000 bp away from both ends of the perfect repeat loci. The dips near x=0 reflect that a non-motif base must precede and follow the perfect region of the repeat at the start and end of the perfectly repeated segment.

### Construction and evaluation of SSR-clouds for detection of SSRs

To characterize and detect oligos in SSR detritus fields, we used the probability clouds (*P-clouds*) method (32, 33), which annotates empirically identified clusters (or clouds) of related oligos that are over-represented in a sequence. This approach has the potential to identify ancient repeats that have diverged considerably from their original sequence. By using increasingly relaxed threshold enrichment parameters, we built nested oligo clouds for each SSR motif family. There are relatively few highly enriched oligos with high similarity to the parent motif, and larger sets of more diverse but less-enriched oligos (Figure 4). High count, high similarity oligos are included in high stringency clouds, and low count, low similarity oligos are built into lower stringency clouds. We note here that although the largest motif families identified over 50,000 16-mer oligos in their low-stringency clouds, this represents only a very small fraction (0.0000116) of all possible 16-mer oligos. We conclude that finding *extended* regions in the genome made up of such oligos by chance alone is improbable. For example, if 50,000 oligos were distributed evenly across the genome, one might expect to find only about one oligo every 100,000 bp.

**Fig. 4.**
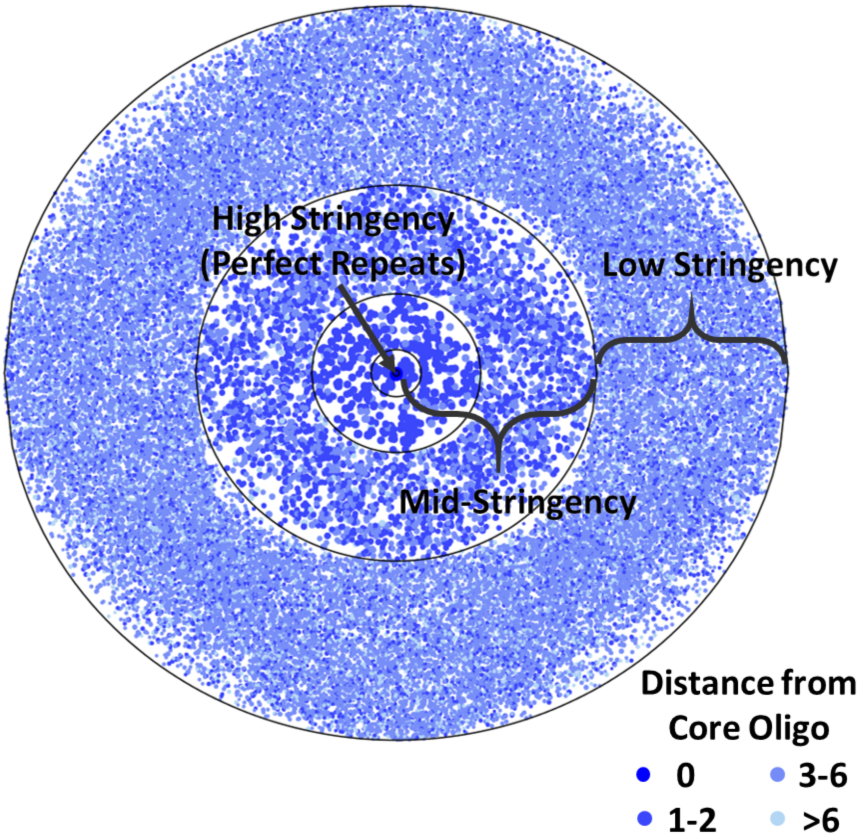
Visual of numbers of poly-A cloud oligos with different similarities from poly-A. Each point represents a 16-mer oligo built into the cloud set for the poly-A SSR family, with oligos clustered into concentric rings depending on distance from the core oligo (poly-A). Darker shades of blue near the center represent higher similarity cloud oligos, and lighter shades represent lower similarity cloud oligos, as indicated in the legend.

SSR-cloud loci were ranked according to the highest-stringency oligo contained in the locus, but annotations of high-stringency oligos can be extended using oligos contained in lower stringency clouds. The extension of locus annotations with lower-stringency oligo clouds has a striking impact on the length distributions of SSR loci (Figure 5). For example, poly-A SSR loci go from a highly skewed, almost exponential length distribution with a mean at 17.2 bp when only perfect repeats are considered, to something much closer to a normal distribution (although still right skewed) with a mean near 36 bp when extended using lower-stringency SSR-cloud sets (Figure 5A). The latter distribution is consistent with the biology of poly-A origins through retrotransposition and reports that *Alu* transposition efficacy increases with poly-A tail length up to 50 bp (42, 43). Thus, the lower-stringency oligos enable detection of a region that is consistent with the *entire* ancient sequence derived from the poly-A tail at the time of insertion. However, it should be recognized that some of the detected length could be due to slippage in either direction post-insertion and prior to degradation. The length distributions of other SSR loci are similarly expanded, but with tails often extending to much larger regions (Figure 5B). Annotation and locus extension may occur infrequently by chance and can be accounted for with false discovery rates. Nevertheless, to ensure that the SSR locus length distributions we observe are not biased towards the loci used in cloud building, we tested the length distributions of the 10% of SSR loci that were not used in cloud building (see Methods). Figure S4 shows that the length distributions of these sets of loci do not substantially change, even at low cloud stringency.

**Fig. 5.**
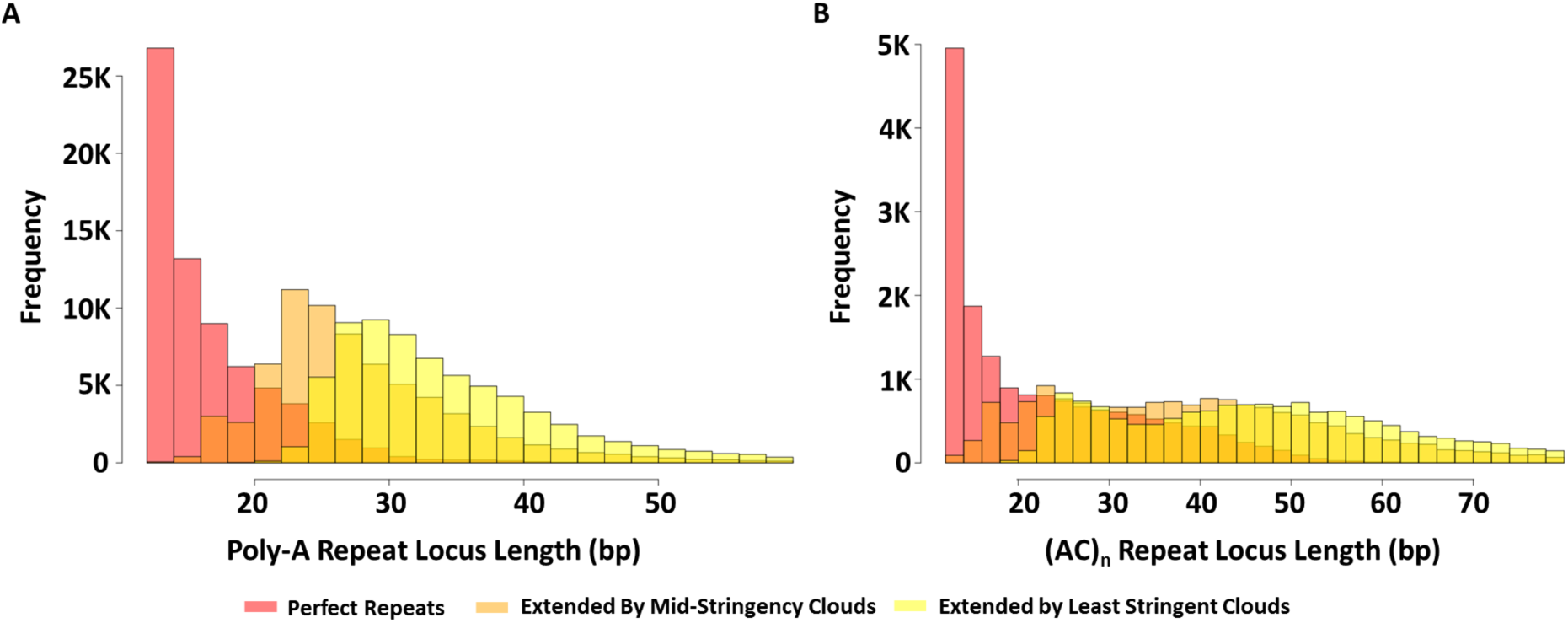
Length distribution of perfect SSR loci annotations expanded using SSR-derived oligos. The length count distributions are shown for: (A) poly-A SSRs; and (B), (AC)_n_ SSRs. Perfect repeat annotations are shown in red, mid-stringency annotations in light orange, and low-stringency annotations in yellow, with darker regions indicating overlap.

### SSR-clouds annotation of the human genome

The complete SSR-clouds annotation comprises 8,983,547 loci covering 221.6 Mb (7.15%) of the human genome. Of these loci, 46.92% intersect a transposable element, which includes poly-A regions annotated as part of the transposable element. A total of 3,085,675 of the loci, comprising 62 Mb (28.15% of all bases annotated by SSR-clouds) do not overlap with any previous repetitive element (including SSRs annotated by TRF), and thus represent novel repetitive sequence. Accounting for false discoveries (see Methods), we conclude that at least 6.77% of the genome is made up of SSRs or is SSR-derived.

The average false discovery rate is 5.31%, but the probability of being a false discovery varies widely among loci, depending on length. Most loci have a high positive predictive value (the inverse of the false discovery rate), but 3,423,735 loci covering 53.8 Mb (∼25% of the SSR-clouds annotation) have a false discovery rate > 10% (maximum FDR= 0.175). The majority (3,020,997, or 88%) of these less certain SSR loci are either 16 bp or 17 bp in length, while the remainder are comprised of short perfect SSR loci under 13 bp in length. Although these loci have high false discovery rates because they are short, there are millions more of these loci than expected by chance based on dinucleotide frequencies. This abundance of short SSRs indicates that simple sequences of this length may often originate during evolution but die quickly through mutation accumulation before they have a chance to extend to create longer loci. It is also worth noting that regardless of their origin, these short loci are identical in sequence to areas that have potentiated SSR expansions and likely good spawning grounds for future SSRs.

### Comparison of SSR-clouds detection to Tandem Repeats Finder

Although the purpose of this research was not to replace Tandem Repeats Finder (TRF), we nevertheless compared the SSR-cloud annotations with TRF annotations using the same parameters as in (2), which yielded the widely-quoted 3% SSR genomic estimation (2). Table 1 (see also Tables S3 and S4) highlights that SSR-clouds annotations of SSRs captures nearly all TRF SSR loci as well as millions of likely SSR-like loci that are not detected by TRF. The greatest increase in SSR-cloud loci occurs where the stringency of the SSR-cloud locus is low. These elements are likely missed by TRF because of their short length or divergence from a perfect SSR sequence. The discordance between the SSR-clouds and TRF annotation sets highlights that previous estimations of SSRs in the genome are likely extremely conservative and frequently overlook SSR-derived regions of more ancient origin. This is conservative in the wrong direction for research questions that require eliminating as many SSR-derived regions as possible, for example if one is trying to identify low-copy regions of the genome or trying to discriminate sequences derived from specific types of TEs, which might themselves include SSRs.

**Table 1.** SSR-clouds recovery of Tandem Repeats Finder (TRF) loci.

|  |  | Highest Cloud Stringency of Locus |  |  |  |  |  | FDR ≤ %5 |  |
| --- | --- | --- | --- | --- | --- | --- | --- | --- | --- |
|  |  | Perfect Repeats |  | Mid-stringency |  | Low Stringency |  |  |  |
|  |  | Loci | bp | Loci | bp | Loci | bp | Loci | bp |
| Poly-A | P-Clouds TRF Intersection | 453,128 | 11,518,426 | 615,893 | 16,085,955 | 556,794 | 17,373,114 | 660,469 | 17,272,038 |
|  | **Total P-Cloud Recovery of TRF** | 67.73% | 62.37% | 92.06% | 87.10% | 83.23% | 94.07% | 98.72% | 93.52% |
|  | Novel Clouds | 244,269 | 13,490,320 | 889,630 | 36,272,378 | 2,282,559 | 65,260,452 | 1,552,401 | 53,363,205 |

Total TRF Poly-A Loci = 669,020 Total TRF Poly-A bp = 18,468,468
|  |  | Highest Cloud Stringency of Locus |  |  |  |  |  | FDR ≤ %5 |  |
| --- | --- | --- | --- | --- | --- | --- | --- | --- | --- |
|  |  | Perfect Repeats |  | Mid-stringency |  | Low Stringency |  |  |  |
|  |  | Loci | bp | Loci | bp | Loci | bp | Loci | bp |
| (AC)<sub>n</sub> | P-Clouds TRF Intersection | 120,498 | 4,813,795 | 143,941 | 5,989,636 | 148,027 | 6,301,466 | 148,027 | 6,301,466 |
|  | **Total P-Cloud Recovery of TRF** | 81.09% | 65.02% | 96.86% | 80.90% | 99.61% | 85.11% | 99.61% | 85.11% |
|  | Novel Clouds | 28,365 | 3,444,295 | 724,496 | 25,393,739 | 1,621,096 | 44,746,021 | 1,621,096 | 44,746,021 |

Total TRF (AC)<sub>n</sub> Loci = 148,607 Total TRF (AC)<sub>n</sub> bp = 7,403,867
|  |  | Highest Cloud Stringency of Locus |  |  |  |  |  | FDR ≤ %5 |  |
| --- | --- | --- | --- | --- | --- | --- | --- | --- | --- |
|  |  | Perfect Repeats |  | Mid-stringency |  | Low Stringency |  |  |  |
|  |  | Loci | bp | Loci | bp | Loci | bp | Loci | bp |
| All SSRs | P-Clouds TRF Intersection | 1,741,873 | 59,642,996 | 1,965,320 | 67,616,136 | 2,119,405 | 71,906,834 | 1,946,410 | 68,221,956 |
|  | **Total P-Cloud Recovery of TRF** | 78.73% | 67.40% | 88.83% | 76.41% | 95.80% | 81.26% | 87.98% | 77.10% |
|  | Novel Clouds | 2,046,914 | 58,749,285 | 2,690,429 | 75,993,192 | 6,702,981 | 149,673,223 | 2,008,354 | 70,732,930 |

Total TRF SSR Loci = 2,212,414
Total TRF SSR bp = 88,485,889
SSR-clouds loci with a merge distance of 5 bp were divided into 3 nested sets based on the most stringent oligo used to annotate each locus. Cells in the table report the number of loci or bp that overlap with TRF loci as well as the number of novel SSR-clouds loci and bp. Comparisons were also made for SSR-clouds loci with FDR ≤ 5%.

### Age characterization of SSR-derived sequences using *Alu* transposable elements

The approximate ages of poly-A SSR-derived sequences were determined by leveraging the relationship between *Alu* transposable elements and poly-A SSRs (15, 40, 44). *Alu* has over a million copies in the human genome, and their relative ages can be accurately determined (37). We divided *Alus* into three age groups approximately representing the main families of *Alu* and assessed how frequently poly-A loci detected by SSR-clouds of different stringencies could be found in the poly-A regions of *Alu* elements. While 63% of young poly-A tails tend to be annotated by uninterrupted poly-A clouds, older poly-A tails from the oldest group of *Alus* (42,125 loci, or ∼50%) are unsurprisingly the most difficult to detect and are often annotated only by low stringency SSR-clouds (Figure 6). These results support the idea that lower-stringency SSR annotations are indeed derived from SSRs but are difficult to detect through other means because of their divergence from the original poly-A repeat.

**Fig. 6.**
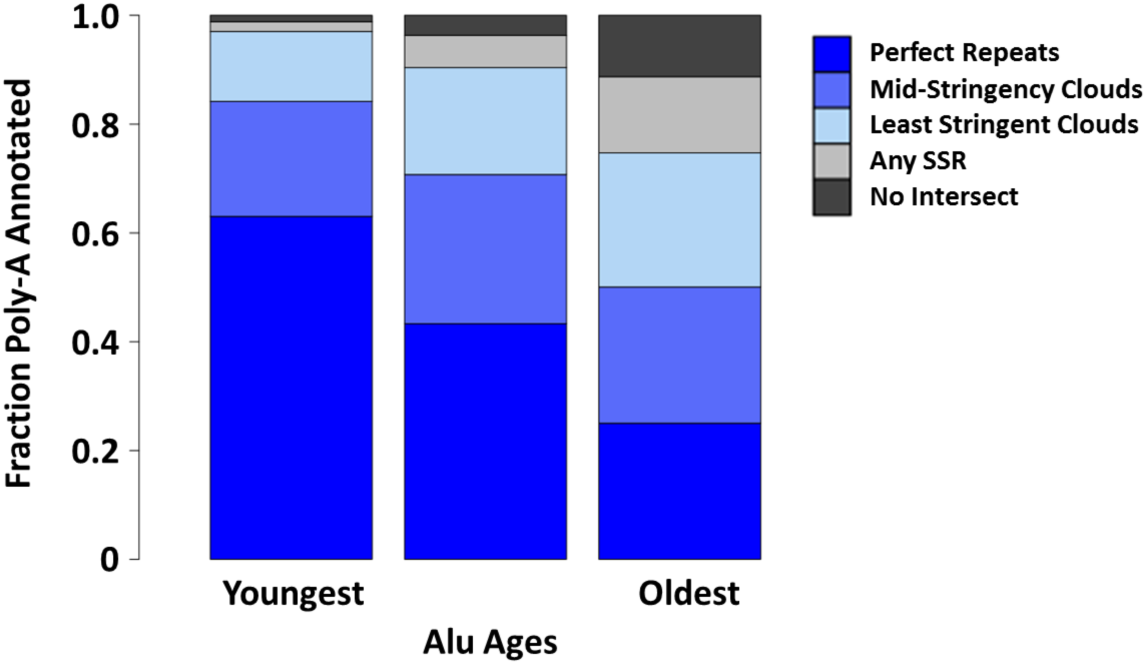
SSR-cloud annotation of poly-A regions adjacent to annotated *Alu*s. Full length *Alus* (275-325 bp) were divided into three groups based on their age (roughly corresponding to the three major expansions of *Alu, AluJ, AluS, and AluY*) and 5’ overlap with poly-A SSR-cloud annotated regions was evaluated. The region expected to carry the poly-A tail was defined as within 30 bp of the *Alu* terminus. Different cloud stringency extensions are colored with dark blue indicating highest stringency poly-A annotations found, and light blue lowest-stringency poly-A annotations. If no poly-A annotations were found, other SSR-cloud loci found are shown in light gray, and no intersecting SSR annotations found shown in dark grey.

About 25% of old loci were not detected by poly-A clouds of any stringency level, but an additional 11,821 annotations were found using SSR-clouds from any SSR family, not just poly-A. Thus, almost 90% of the oldest *Alus* (74,846 loci out of 84,346 total) had some sort of SSR-derived locus in the expected poly-A region. It is possible that the 9,500 old *Alus* without detected SSR-clouds had their tails deleted or moved through genomic rearrangements over time or they degenerated to the point of being unidentifiable. The oldest group of *Alus* is 1.60 times older than the average age for all *Alus*, while the unannotated *Alus* are 1.64 times older (Welch two-sample t-test, p < 2.2 x 10^-16^), supporting the idea that loss of tails increases with age.

## Discussion

SSR-clouds is a rapid, non-parametric method based on *P-clouds* for finding SSRs and SSR-derived regions in the genome. SSR-clouds finds numerous previously undiscovered SSR loci whose overlap with poly-A regions of known ancient transposable element loci provides compelling evidence that these loci are indeed SSRs or are SSR-derived. SSR-clouds analyses reveal that SSR-derived regions comprise a larger portion of the human genome than previously appreciated, increasing the SSR-derived percentage from about 3% to at least 6.77%. This increase is due to increased annotation length of previously annotated loci as well as newly annotated loci (Table 1). The output for SSR-clouds follows a standard bed file format (including the chromosome/scaffold and beginning and ending coordinates for a locus), with additional information about the SSR motif family present in the locus. As seen in Figure 7, different regions of a locus may be annotated by the clouds of multiple families, creating a complex locus. For complex loci, SSR-clouds gives information about each of the families present in the locus, including the average cloud stringency of that family’s oligos in the locus and what percentage of the locus is covered by oligos from that family’s clouds. We consider this output, which simultaneously considers all families that may be present in a locus, to more accurately reflect the true nature of SSRs, given the propensity of SSRs to spawn different SSR motif families during their evolution.

**Fig. 7.**
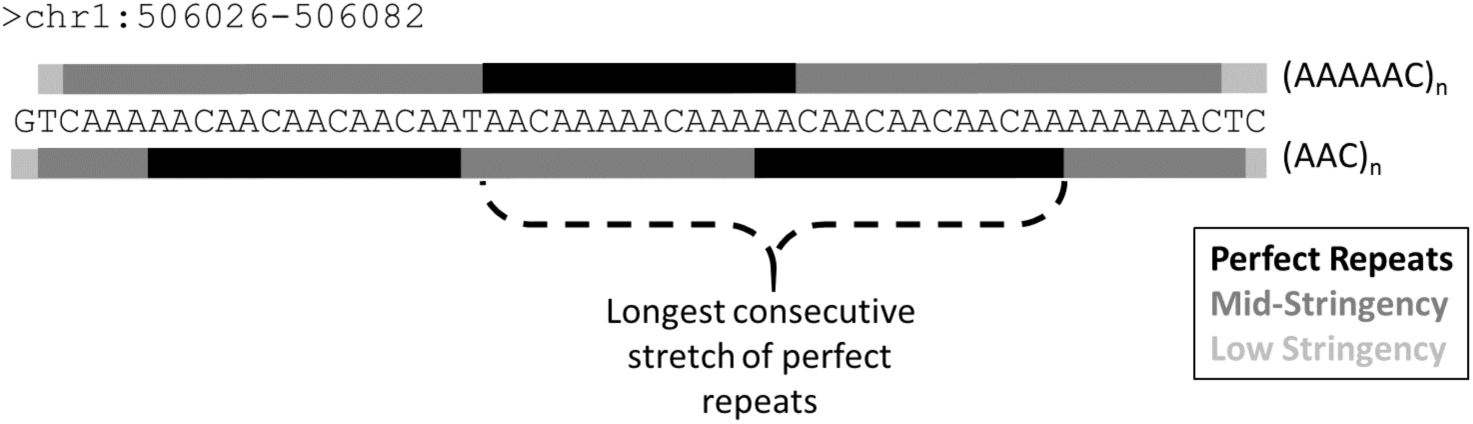
Anatomy of a complex SSR locus and its annotation by SSR-clouds. The sequence for an SSR locus found at bp 506026-506082 on chromosome 1 in hg38 is shown. Regions annotated by the two most prevalent families, AAAAAC (top) and AAC (bottom), are shown, with perfect repeats indicated with a black bar, mid-stringency cloud annotations with a dark gray bar, and the lowest stringency cloud annotations with a light gray bar. The longest stretch of perfect repeats of any kind (26 bp) is indicated, and was used to determine the false discovery rate of the locus (see Methods).

By identifying millions of previously overlooked short and imperfect SSR loci, we provide evidence that the SSR life cycle is highly flexible and show that multiple paths to SSR death exist. While some of the short loci may be fossils of longer ancient loci that are no longer detectable, our analysis of *Alu* poly-A’s suggests that only ∼10% of mature SSR loci fall below detectability even after 65 million years. It thus seems reasonable that a substantial fraction of these short loci are more frequent than expected from point mutation processes and therefore created by some amount of slippage, but never reached SSR maturity where slippage events would have rapidly increased the locus size, and instead died in their infancy. Regardless of their precise origins, it is reasonable to think that these short loci may yet act as birthing grounds and nurseries for future SSRs, thus creating another alternate route through the SSR life cycle without ever passing through adulthood. The abundance of these short SSR-derived loci also indicates that SSRs may be born much more frequently than appreciated; with nearly 9 million separate loci, there is an average of one SSR for every 350 bp.

An important feature included in SSR-clouds that is lacking in standard SSR annotation software is the estimation of false discovery rates for each locus. Recently active SSR loci can be identified with high confidence because they have spent little time in the genomic churn caused by mutation and fragmentation, but this is not the case for millions of ancient SSR loci that we identified here. We note that even the short loci with high false discovery rates, although they may not be derived from mature SSR loci with high slippage rates, may be important to identify as potential sources of new SSR loci. Furthermore, loci with high false discovery rates can be included or excluded in downstream analyses based on user-defined analysis-specific false discovery thresholds and the needs and tolerances of the researchers for both false discoveries and failure to detect relevant elements. Figure S5 illustrates the effect of different false discovery thresholds on the total number of base pairs identified as SSRs in the human genome.

## Conclusions

We extend previous reports of sequence bias near SSR loci (29, 30) and show that the boundaries of this bias, though motif dependent, may extend for hundreds of base pairs to either side of an SSR locus. The length of sequence bias near SSR loci indicates that distinct boundaries on the distance of SSR spawning events exist, and the data presented here suggests that such events are generally limited to within several hundred base pairs of parent loci. Our characterization of similarity between clustered SSR loci supports this assertion and provides further evidence that the generation of new SSR loci is greatly influenced by the evolution of locally active SSRs.

Because the motif, purity, and length-dependent nature of SSR locus evolution is complex, the SSR-clouds approach presents an important and tractable method to improve studies of the different phases of the SSR life cycle that cannot be easily achieved through other approaches. The data presented here reveal unprecedented detail into the proposed SSR life cycle (15–18). The signals of highly biased sequence near SSR loci and clustered similar loci (see Figs 1-3) can be generated through repeated rounds of interrupting mutations within an SSR locus to isolate regions of the locus followed by expansion in regions that remain susceptible to slippage. This process of constant sloughing off of SSR detritus can be likened to simultaneous birth and death processes, and creates natural boundaries at SSR loci, which we report here. This process also makes predictions about SSR sequence degeneracy over time possible; long dead SSR loci resemble the derived and most degenerate portions of active SSR loci that are near the boundaries of the SSR locus.

A large fraction of recent (4-6 million years old) *Alu* elements (∼60%) have intact poly-A tails, and only a small fraction (<5%) have different motifs or no SSR at all in their poly-A tail region. Notably, the remaining nearly 40% have already begun to degenerate, even after relatively recent successful retrotransposition. However, although the poly-A appears to rapidly degenerate, these degenerate regions are detectable in many of even the oldest of *Alu* elements, demonstrating both a surprising longevity of SSR character in ancient simple repeats, and the sensitivity of SSR-clouds method.

The longevity of SSR loci is further highlighted by the fact that a substantial proportion (∼15%) of poly-A’s from the oldest group of *Alus* spawned new SSRs with different motifs (Figure 6). Spawning of SSRs has not been characterized in great detail (15), but this evidence, combined with the tendency of similar SSR repeats to cluster, presents a timeline for spawning events while also characterizing the expected motif bias for newly spawned loci.

The high degree of overlap between transposable elements and SSR loci we present here supports the hypothesis that transposable elements play a substantial role in the generation of SSR loci (27, 40, 44). About half (46.92%) of SSRs intersect with an easily-identifiable transposable element. Because about half the genome is made up of easily-identifiable transposable elements (1), this might suggest that SSR origins are similar in TE and non-TE regions. However, evidence suggests that many transposable elements in the ‘dark matter’ portion of the genome are not-so-easily-identifiable (32, 33). It seems likely that a large fraction of the remaining SSRs were generated through the action of the hard-to-identify old and fragmented elements. Due to the ability of an SSR locus to maintain SSR character over long periods of time through constant slippage and spawning, the SSR loci identified by SSR-clouds may yet provide additional information in identifying the origins of ‘dark matter’ in the genome.

## Methods

### Annotation of perfect SSRs and surrounding regions

Oligonucleotide sequences representing all possible SSR sequences were created *in silico* using a *Perl* script that clusters alternate phases of the same SSR motif (ACT = CTA = TAC) and reverse complements of each phase into a single motif family. Perfect SSR repeat loci were defined as uninterrupted tandem repeats of a single motif family ≥ 12bp in length, and perfect stretches separated by 1 bp or more non-motif nucleotides were considered different loci. Perfect SSRs, as defined above, were annotated in an unmasked version of hg38.To identify sequence bias in regions near perfect SSR loci, each kmer *(k*-length oligonucleotide sequence) within 1,000 bp of a perfect repeat locus was compared with the kmers from different phases of the perfect motif. Mean similarities to the closest repeat kmer were calculated versus distance from locus boundaries, and distances between perfect SSR repeat loci were also recorded.

### Constructing SSR-clouds

SSR-clouds were constructed similarly to cloud construction methods outlined in (32, 33) with modifications described here. To construct p-clouds from SSR-flanking regions we conservatively used 16-mer oligonucleotides and considered only 50 bp on either side of a perfect repeat locus as a template for cloud formation. P-clouds for each SSR motif family were constructed separately from one another using a training set that consisted of a randomly chosen subset of 90% of loci for each family, with the remaining 10% of loci used as annotation tests. Loci that were separated by fewer than 100 bp from other loci of the same family were merged into a single locus before cloud formation to prevent double counting oligos in the regions between the loci. Following standard *P-cloud*s formation protocol (32), p-clouds were organized around 16-mer core oligonucleotides, including every 16-mer oligo with count above the threshold that was within one nucleotide of the cloud core or any other oligo already in a cloud. For each motif family, we created nested oligonucleotide clouds using lower threshold counts for clouds of lower stringency, such that all oligonucleotides of higher stringency clouds were included in lower stringency clouds. Perfectly repeated 12-mer oligonucleotides were also automatically added to the highest stringency cloud. Different threshold counts were used as criteria for inclusion in p-cloud sets for each motif family depending on the total number of perfect loci used for cloud training, though motif families with fewer than 100 loci in the training set were not used in cloud building. These thresholds, the number of loci used in cloud formation, and the counts of unique oligonucleotides in each stringency level are specified in Table S1. Transposable elements (e.g., *Alu* in humans) were not our targets but are highly represented in regions flanking SSRs, and so all transposable elements annotated by RepeatMasker (35) were removed prior to cloud formation. Because clouds were formed separately for each family, individual oligonucleotides, including those representing perfect repeats, can belong to cloud sets for multiple families.

Annotation with SSR-clouds was performed in an unmasked version of hg38 by simultaneously mapping oligonucleotide clouds from all motif families, and then merging loci within 5 bp of each other into a single locus. Annotations with merge distances of 0 bp and 30 bp were also performed and are presented as supplements. After annotation, loci were ranked and separated according to the highest stringency cloud found in the locus. In analyses presented here that use only single motif families, (poly-A and (AC)_n_), annotation was performed in the same way except that only oligonucleotides created from that family were used.

### Simulating genomes to obtain false positive rates

Fifteen simulated genomes were created from the human genome (hg38) using nucleotide and dinucleotide frequencies obtained from1 Mb windows along the genome. Prior to creation of the simulated genomes, all regions annotated as either a perfect SSR or annotated as transposable elements or other repeat regions by RepeatMasker were masked so that they would be representative of non-repetitive portions of the genome. The simulations proceeded by randomly selecting nucleotides conditional on the dinucleotide frequencies. When the previous nucleotide was absent or undetermined, a starting nucleotide was selected based on independent single nucleotide frequencies. SSR clouds were annotated in the simulated genomes exactly as done for the actual genome. False positive rates for each locus length (or longer) were calculated, for each cloud stringency setting, as the cumulative amount of simulated sequence annotated, divided by the amount of sequence analyzed. Under a given stringency setting, the length of a locus was considered to be the longest stretch of the locus that was consecutively annotated. False positive rates for each locus length and cloud stringency category were calculated for hg38. False discovery rates were then calculated as the expected cumulative falsely annotated sequence, conservatively assuming the entire genome is not SSR, divided by the observed cumulative length annotated for each setting

### Comparison with Tandem Repeats Finder Annotations

Tandem Repeats Finder (TRF) (31) version 4.07b was run under the two parameter sets described in Warren et al. 2008 that were applied to the human genome (hg38) with centromeres and telomeres masked. The two resulting annotation sets were merged to obtain the TRF annotation used here. TRF SSR annotations were segregated into groups by motif family and annotations within each family were merged using BEDTools (36). The BEDTools Intersect function was used to search for SSR-clouds annotations that overlapped with TRF SSR annotations and to determine the number of novel SSR-clouds annotations.

### Intersection with Poly-A Regions of Alu Elements for Age Analysis

Full-length and non-concatenated *Alu* elements were obtained by filtering RepeatMasker *Alu* annotations from the hg38 assembly of the human genome. Relative ages of each element (measured in inferred number of substitutions since retrotransposition) were then estimated by applying the AnTE method to this dataset (37). We began with 823,789 individual full-length *Alu* elements, with each element having an estimated age or retrotransposition relative to the mean age of retrotransposition of all *Alu* elements. To maximize the chances that the *Alus* tested still contained their poly-A tail, we removed all *Alus* that were < 275 bp or > 325 bp in length as well as those *Alus* that were within 50 bp of another TE. After filtering, 407,438 *Alus* remained.

The remaining *Alu* annotations were split into three groups by age and roughly based on the major expansions of *AluY*, *AluS*, and *AluJ*. The youngest group consisted of 57,873 *Alu* elements, ∼97% of which are classified as *AluY* by RepeatMasker, with a mean age of 0.51 relative to the mean age of all *Alus*. The second and largest group, 99% of which are classified as *AluS* elements, consisted of 265,219 elements with a mean age of 0.92 relative to the mean age of all *Alus*. The third group consisted of all *Alu* elements older than those included in the first two groups, 90% of which are classified as *AluJ* and 10% as *AluS*, and had 84,346 elements with a mean age of 1.6 relative to the mean age of all *Alus*.

To ensure detection of only the poly-A region of *Alu* rather than other SSR-rich regions in *Alu*, we used the 30 bp directly 3’ to each *Alu* tested for intersection. We used the *intersect* function from BEDTools (36) to count the number of *Alu* elements that intersected each of the poly-A SSR annotations, beginning with the highest stringency poly-A annotations and proceeding to the lowest stringency annotations.

## Declarations

### Availability of data and materials

The SSR-clouds software and datasets generated and/or analyzed during the current study are available either in the GitHub repository, https://github.com/popgengent/SSRclouds, or from the corresponding author on reasonable request. The P-clouds SSR package was written and implemented in *Perl*. The program parameters can be easily modified for different applications and sensitivity via either the command line or a control file.

### Competing interests

The authors declare that they have no competing interests.

### Funding

This work was supported by the National Institutes of Health (NIH; GM083127 and GM097251 to DDP).

### Authors’ contributions

JAS developed code, designed, performed, and interpreted analyses, and was a major contributor in writing the manuscript; RPR contributed code, designed, and interpreted analyses; CC developed *Perl* version of *P-clouds* used in SSR-clouds code; ACW contributed the library of *Alu* elements and determined their times of retrotransposition; DDP designed, consulted, and interpreted analyses, and was a major contributor in writing the manuscript.

## Acknowledgements

Not applicable.

## Supplemental Figure and Table Legends

**Fig. S1.**
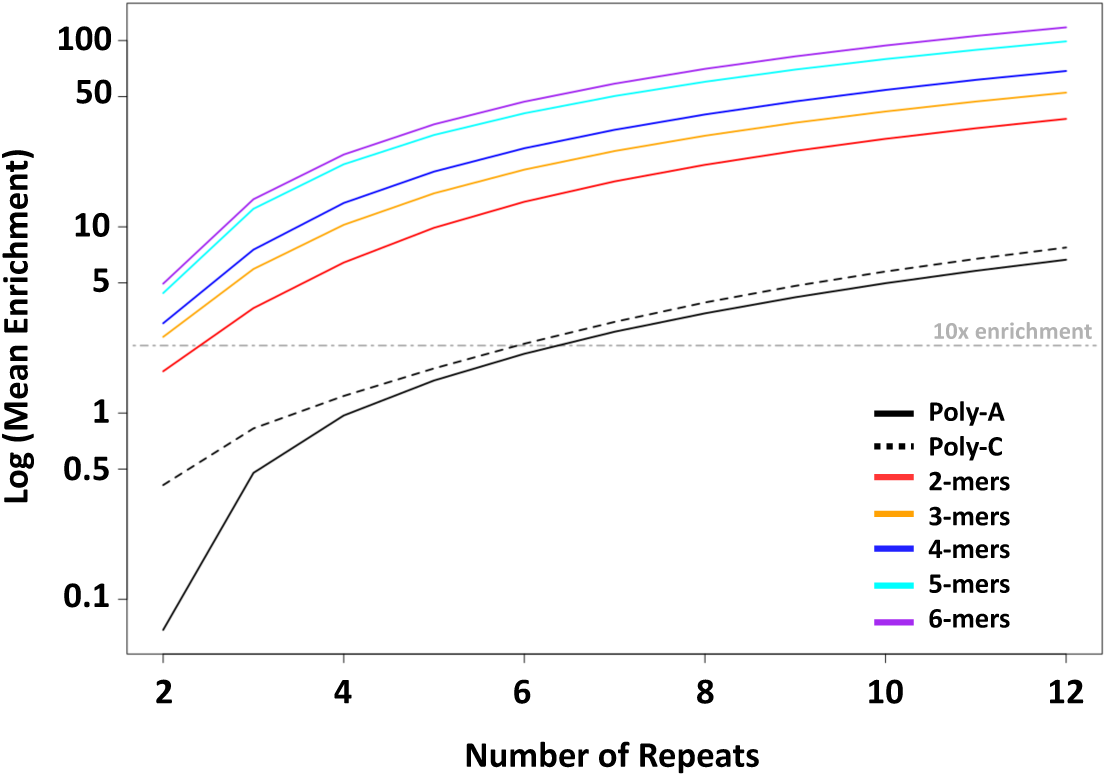
Enrichment of SSRs in the human genome. The mean enrichment of perfect repeats is shown relative to expectation from single nucleotide frequencies. All SSR motifs of a given length were clustered into groups, except that the Poly-A and poly-C single nucleotide repeats are shown as separate lines. The enrichment is shown for the number of repeats of a given size observed in tandem, and the gray dashed lines indicate 10x, 100x, and 1000x enrichments.

**Fig. S2.**
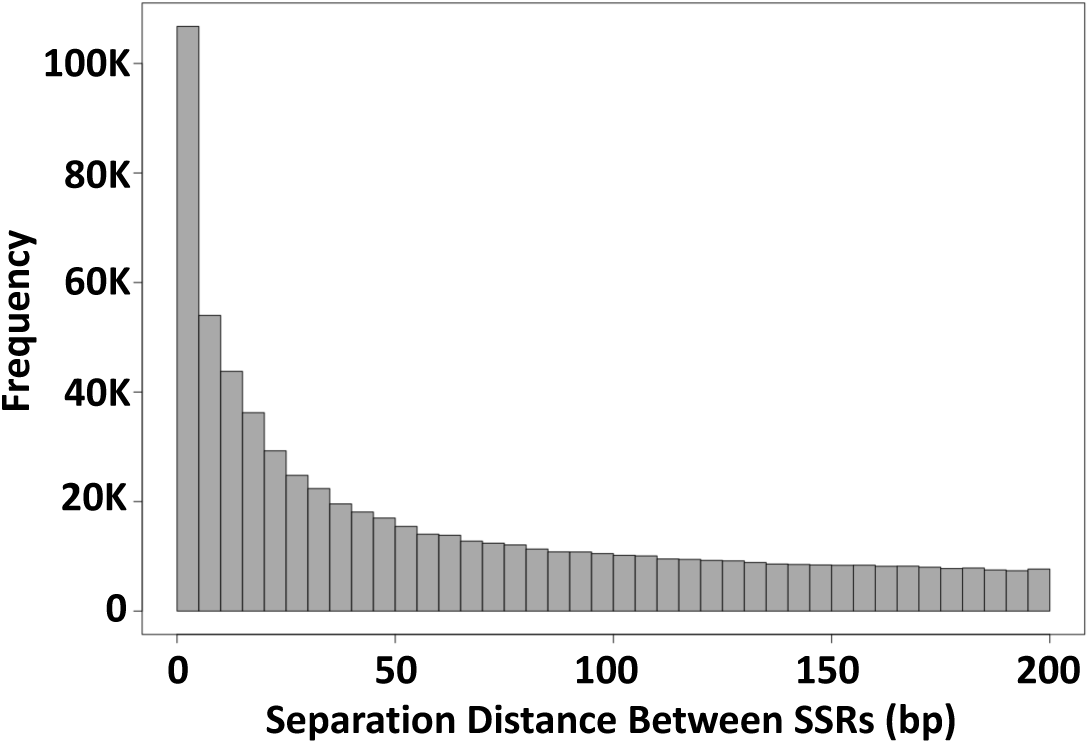
Separation distance between perfect SSRs in the human genome. The frequency of pairs of perfect SSRs ≥ 12 bp long with a given separation distance is shown. The separation distances were binned into groups of 5. The results in A) are for a masked version of the human genome, while B) shows results for an unmasked genome, demonstrating the strong effect and particular features of transposable element SSRs.

**Fig. S3.**
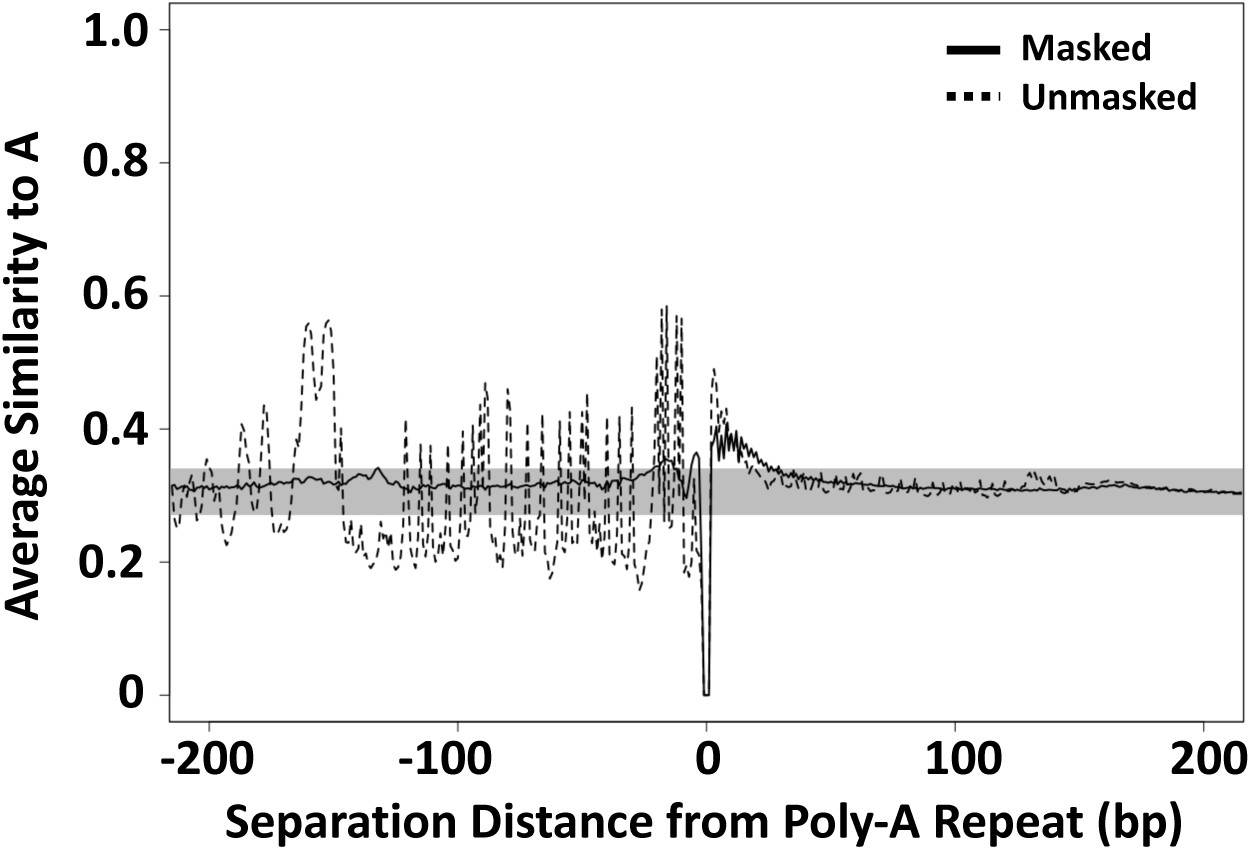
Asymmetric similarity to poly-A. The frequency of adenine nucleotides (A) at every site within 200 bp of perfect poly-A repeats. The solid line shows the frequency of A in a human genome where all transposable elements have been masked and the dotted line shows the frequency in an unmasked human genome. As a reference, the gray box represents a range of 3 standard deviations from the mean frequencies of A calculated in 700 bp windows from 300-1,000 bp away from both ends of all perfect repeats. The strongly varying frequencies in the unmasked genome are mostly a symptom of the high copy number of retroelements such as Alu and Line1. The asymmetric frequency of A’s adjacent to perfect A repeats in the masked genome likely reflects incomplete masking of transposable elements and the existence of other unmasked retrotransposed sequences in what would have been the 5’ region of the retrotransposed poly-A mRNAs.

**Fig. S4.**
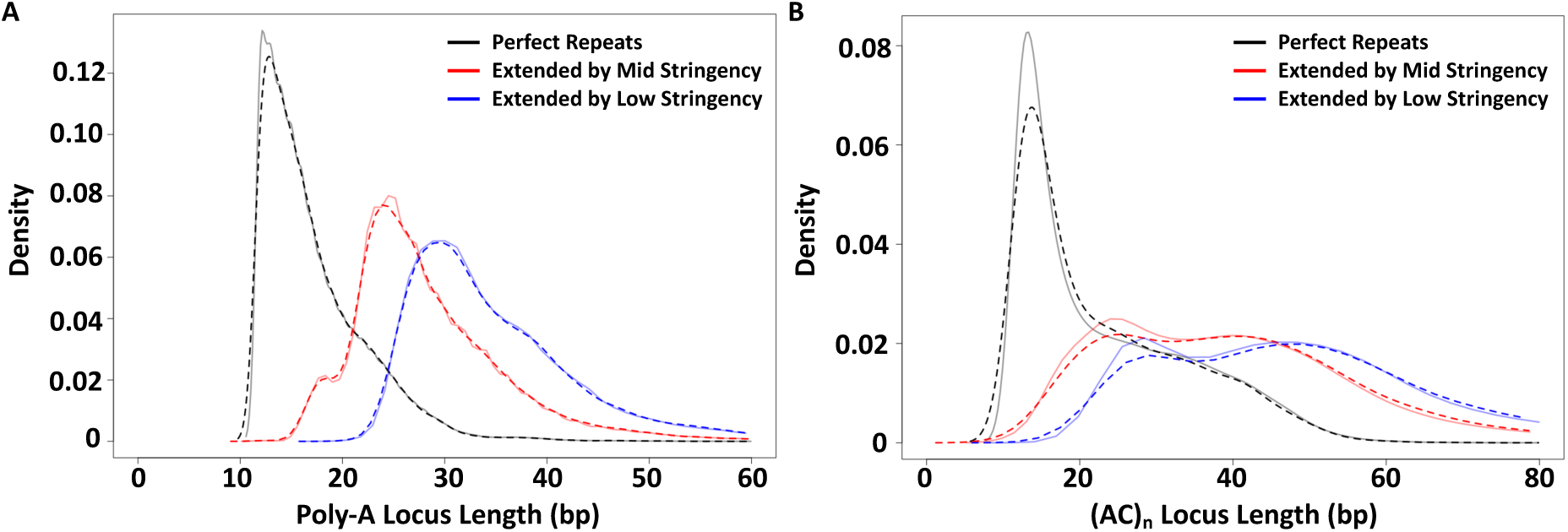
Cloud extension length distributions of training and test loci. Locus length density plots of SSR loci containing perfect repeats (black) and lengths after extension by mid-(red) and low-stringency (blue) cloud sets. Solid lines depict the distributions of lengths for training loci and dashed lines depict the almost perfectly overlapping distributions of lengths for test loci.

**Fig. S5.**
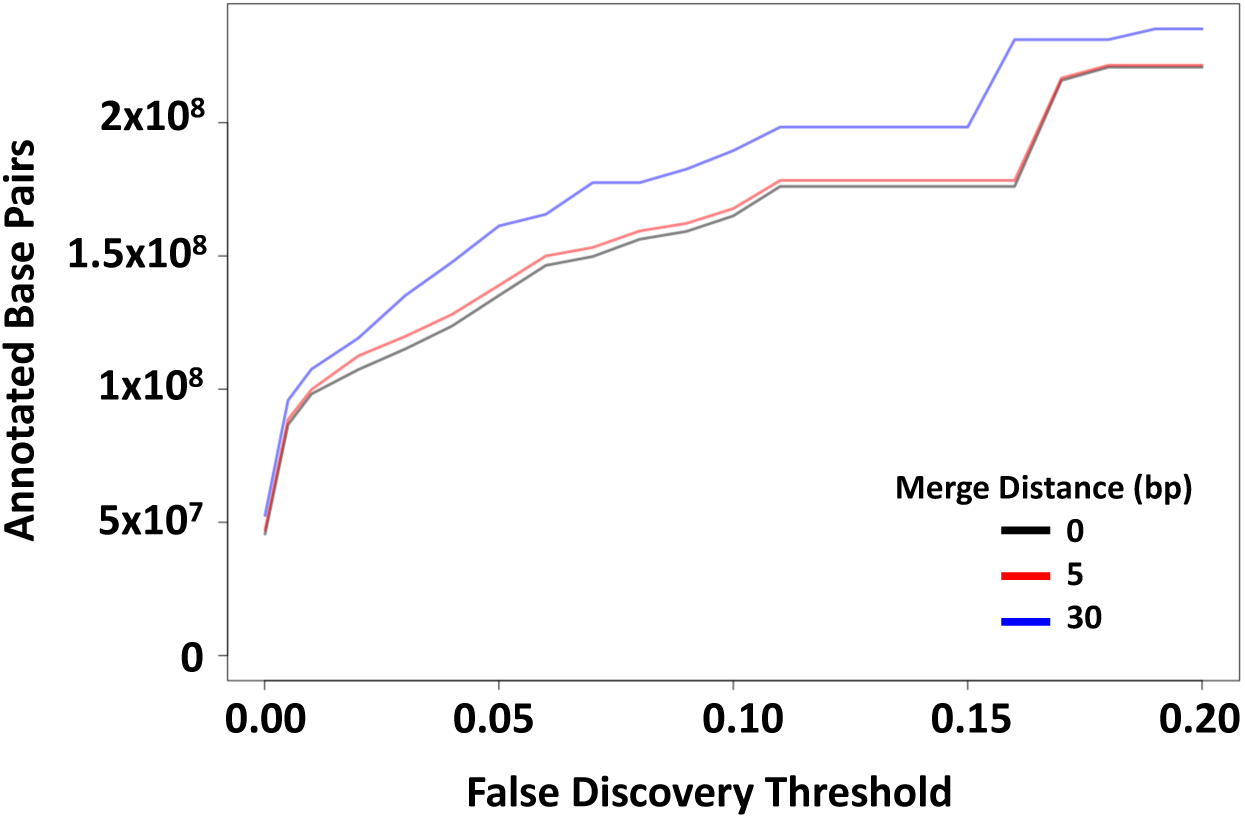
Genomic SSR content annotated with different merge distances and false discovery thresholds. The number of bp in the human genome that were annotated by SSR-clouds under various conditions are shown. with different merge distances and false discovery thresholds. Three lines are shown for merge distances of 0 bp (black), 5 bp (red), and 30 bp (blue), with the per-locus maximum false discovery criterion on the X axis.

Tables S1 and S2 are included at the very end of the manuscript due to their length

**Table S3.**
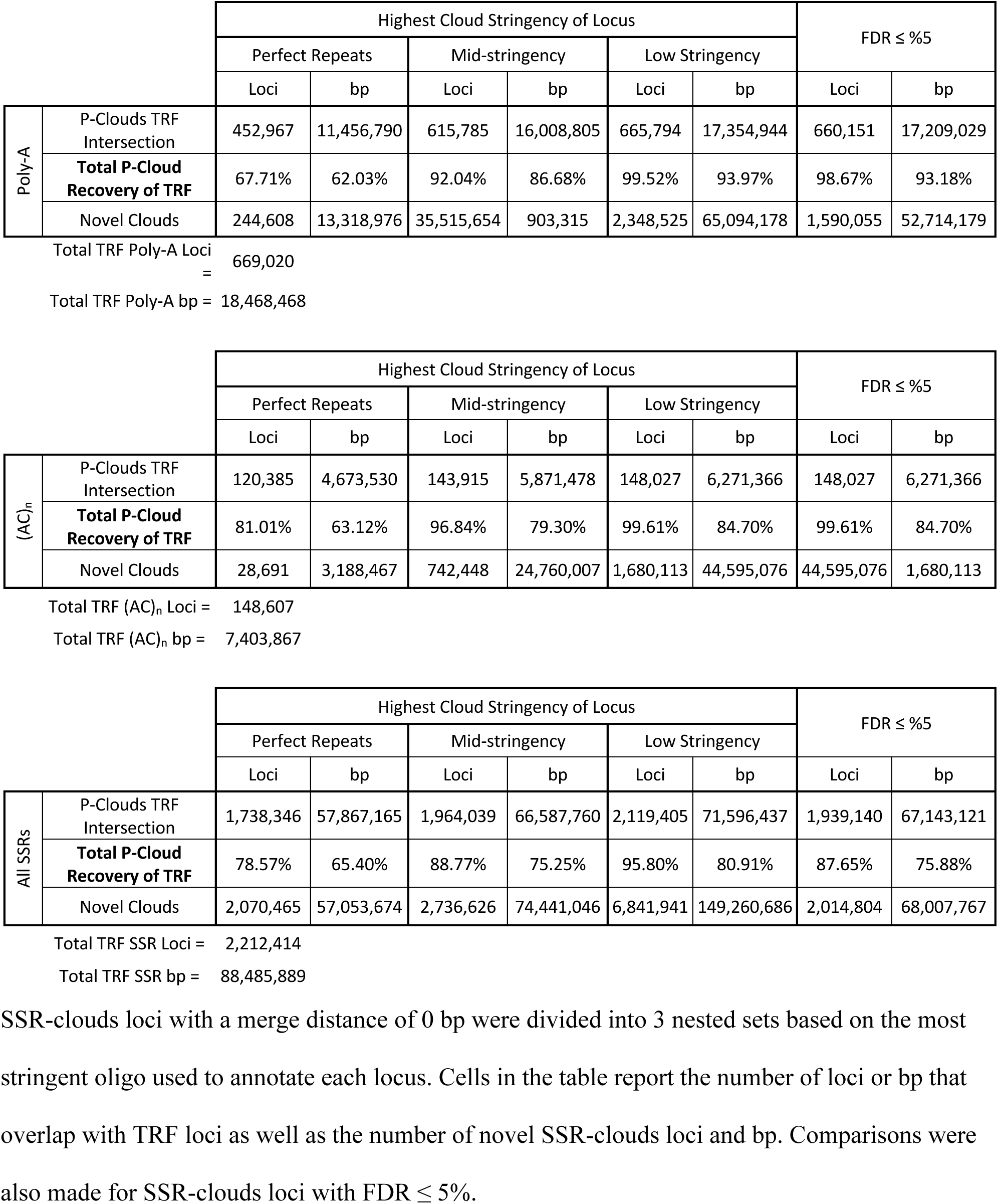
SSR-clouds recovery of Tandem Repeats Finder (TRF) loci.

**Table S4.** SSR-clouds recovery of Tandem Repeats Finder (TRF) loci.

|  |  | Highest Cloud Stringency of Locus |  |  |  |  |  | FDR ≤ %5 |  |
| --- | --- | --- | --- | --- | --- | --- | --- | --- | --- |
|  |  | Perfect Repeats |  | Mid-stringency |  | Low Stringency |  |  |  |
|  |  | Loci | bp | Loci | bp | Loci | bp | Loci | bp |
| Poly-A | P-Clouds TRF Intersection | 454,655 | 11,649,451 | 616,750 | 16,209,662 | 665,794 | 17,446,377 | 660,870 | 17,373,267 |
|  | **Total P-Cloud Recovery of TRF** | 67.96% | 63.08% | 92.19% | 87.77% | 99.52% | 94.47% | 98.78% | 94.07% |
|  | Novel Clouds | 241,932 | 14,420,787 | 862,842 | 39,084,443 | 2,151,156 | 67,376,882 | 1,484,085 | 56,519,523 |

Total TRF Poly-A Loci = 669,020 Total TRF Poly-A bp = 18,468,468
|  |  | Highest Cloud Stringency of Locus |  |  |  |  |  | FDR ≤ %5 |  |
| --- | --- | --- | --- | --- | --- | --- | --- | --- | --- |
|  |  | Perfect Repeats |  | Mid-stringency |  | Low Stringency |  |  |  |
|  |  | Loci | bp | Loci | bp | Loci | bp | Loci | bp |
| (AC)<sub>n</sub> | P-Clouds TRF Intersection | 120,822 | 5,311,956 | 144,057 | 6,394,349 | 148,027 | 6,573,040 | 148,027 | 6,573,040 |
|  | **Total P-Cloud Recovery of TRF** | 81.30% | 71.75% | 96.94% | 86.36% | 99.61% | 88.78% | 99.61% | 88.78% |
|  | Novel Clouds | 27,486 | 4,207,203 | 696,337 | 27,533,146 | 1,524,044 | 46,227,577 | 1,524,044 | 46,227,577 |

Total TRF (AC)<sub>n</sub> Loci = 148,607 Total TRF (AC)<sub>n</sub> bp = 7,403,867
|  |  | Highest Cloud Stringency of Locus |  |  |  |  |  | FDR ≤ %5 |  |
| --- | --- | --- | --- | --- | --- | --- | --- | --- | --- |
|  |  | Perfect Repeats |  | Mid-stringency |  | Low Stringency |  |  |  |
|  |  | Loci | bp | Loci | bp | Loci | bp | Loci | bp |
| All SSRs | P-Clouds TRF Intersection | 1,762,767 | 64,154,932 | 1,974,700 | 70,907,679 | 2,119,427 | 74,288,802 | 1,956,493 | 71,532,092 |
|  | **Total P-Cloud Recovery of TRF** | 79.68% | 72.50% | 89.26% | 80.13% | 95.80% | 83.96% | 88.43% | 80.84% |
|  | Novel Clouds | 1,922,645 | 73,344,350 | 2,504,656 | 91,070,015 | 6,072,473 | 160,927,330 | 1,889,528 | 89,725,301 |

Total TRF SSR Loci = 2,212,414
Total TRF SSR bp = 88,485,889
SSR-clouds loci with a merge distance of 30 bp were divided into 3 nested sets based on the most stringent oligo used to annotate each locus. Cells in the table report the number of loci or bp that overlap with TRF loci as well as the number of novel SSR-clouds loci and bp.
Comparisons were also made for SSR-clouds loci with $FDR \leq 5\%$ .

**Table S1.**
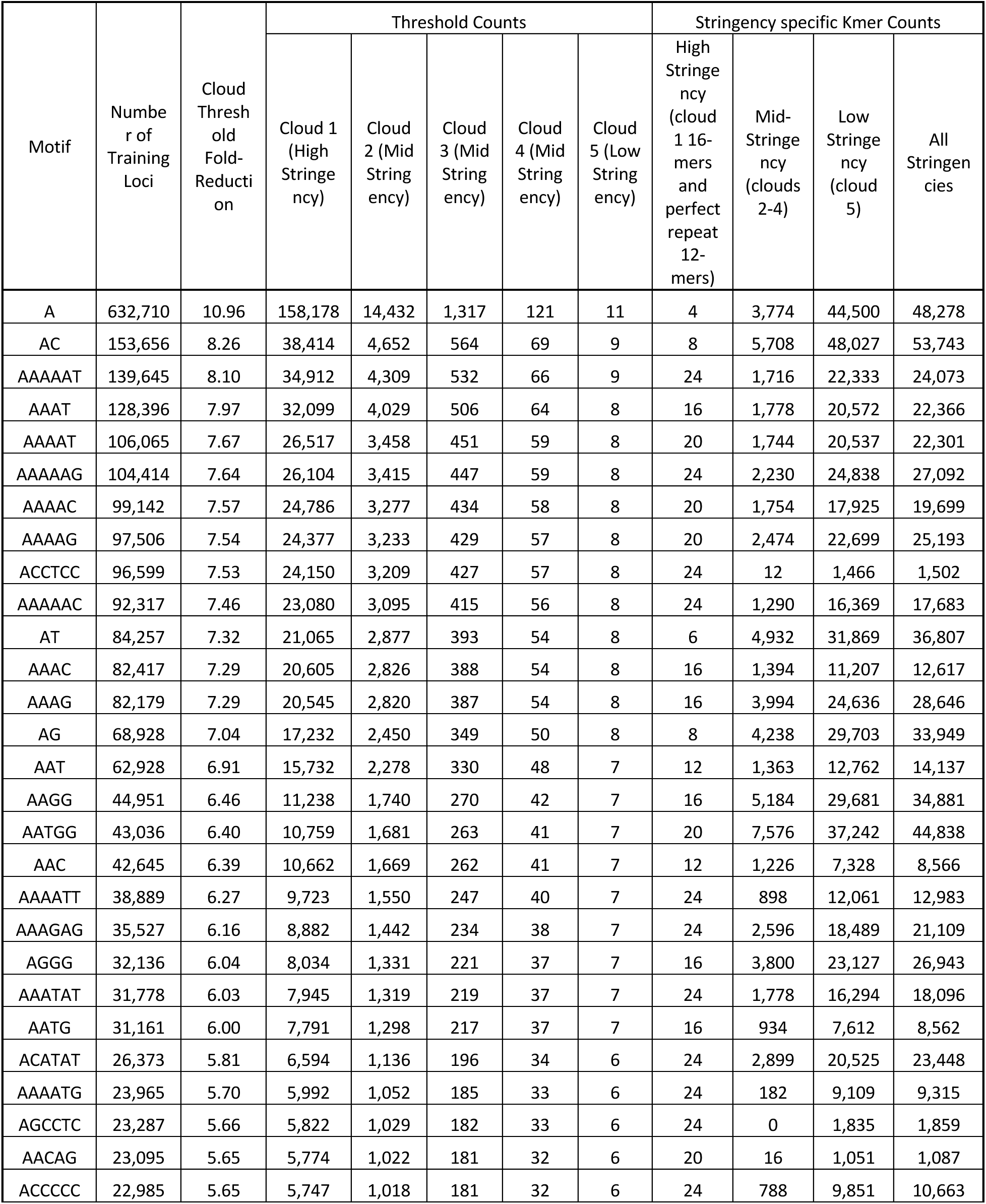
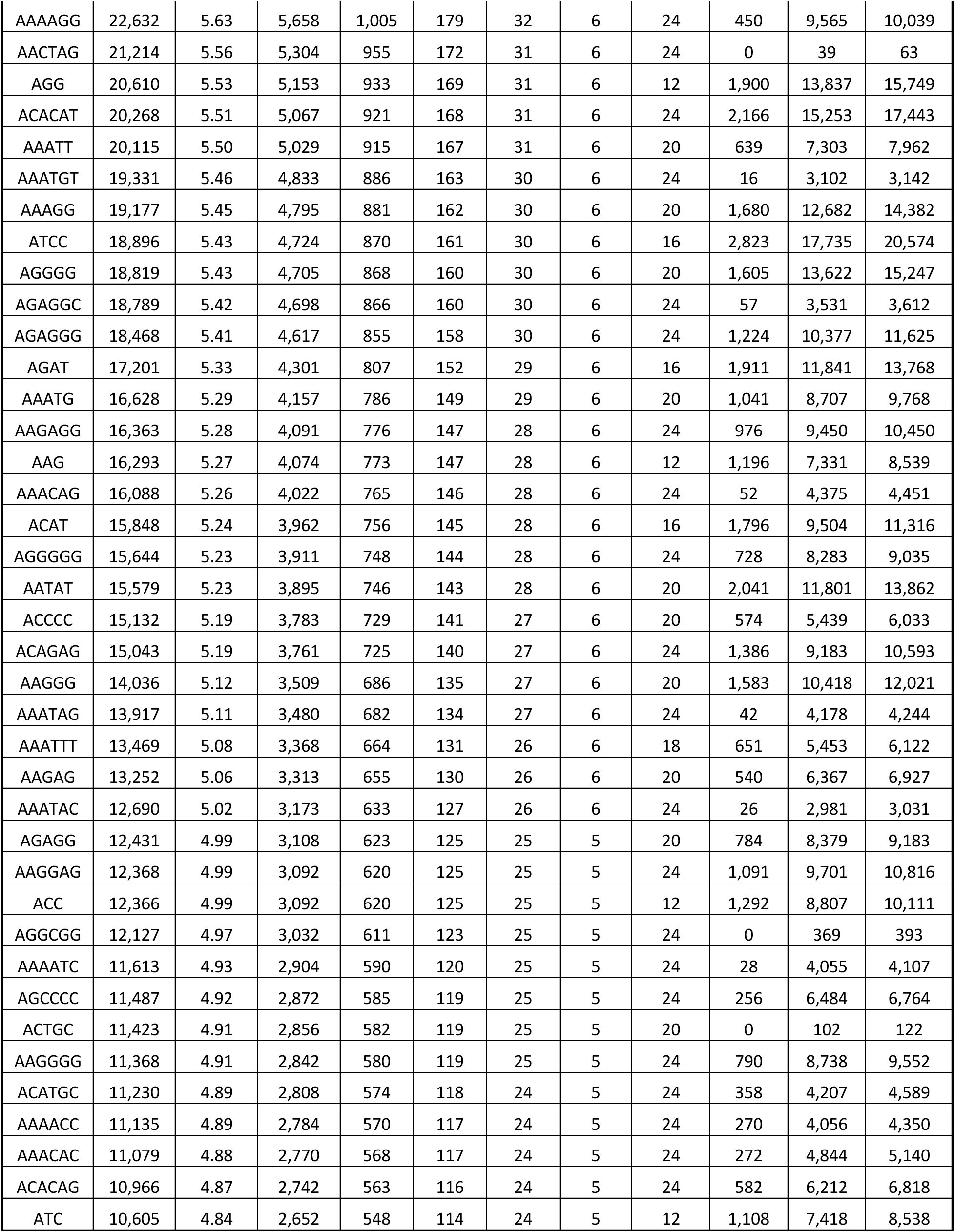
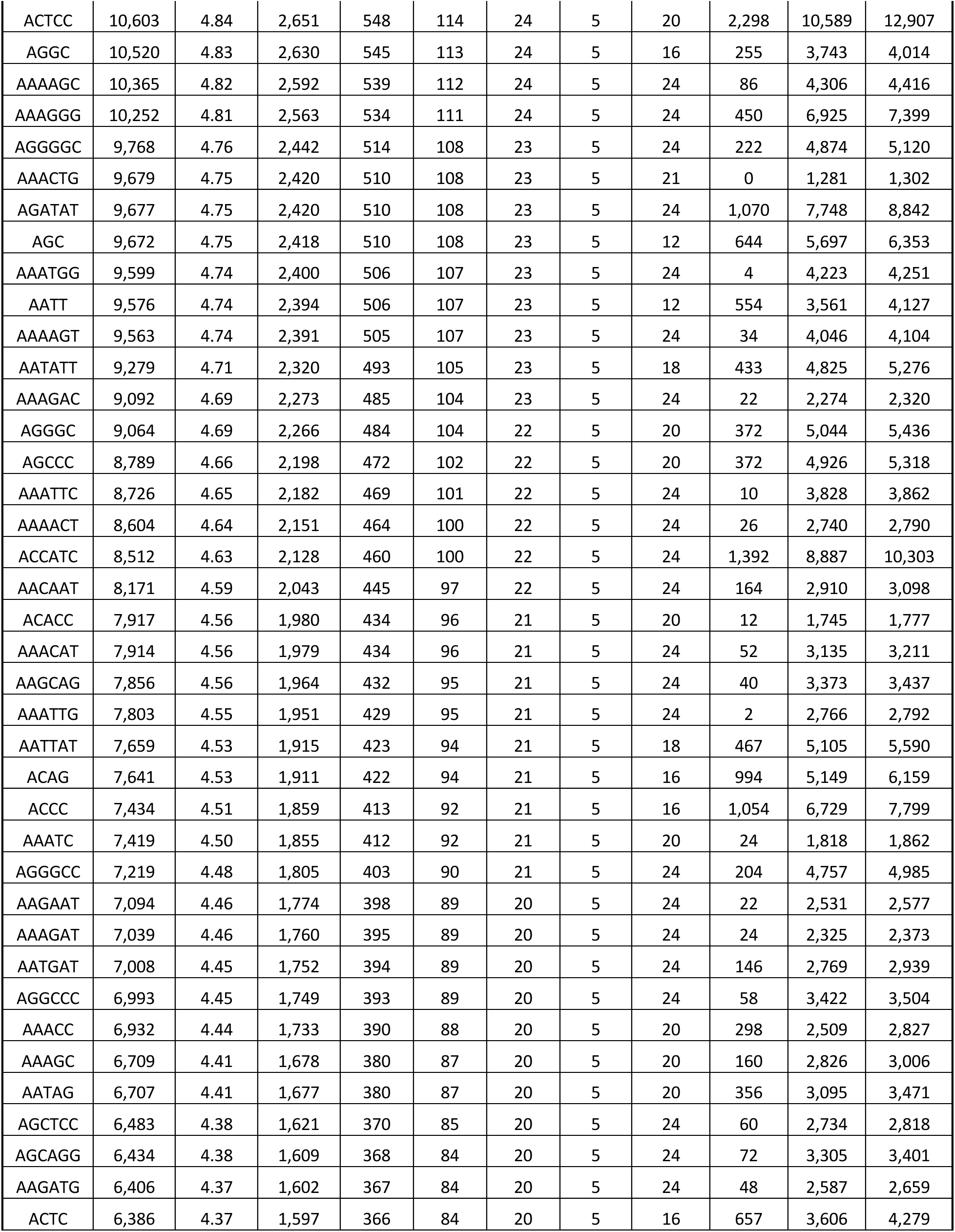
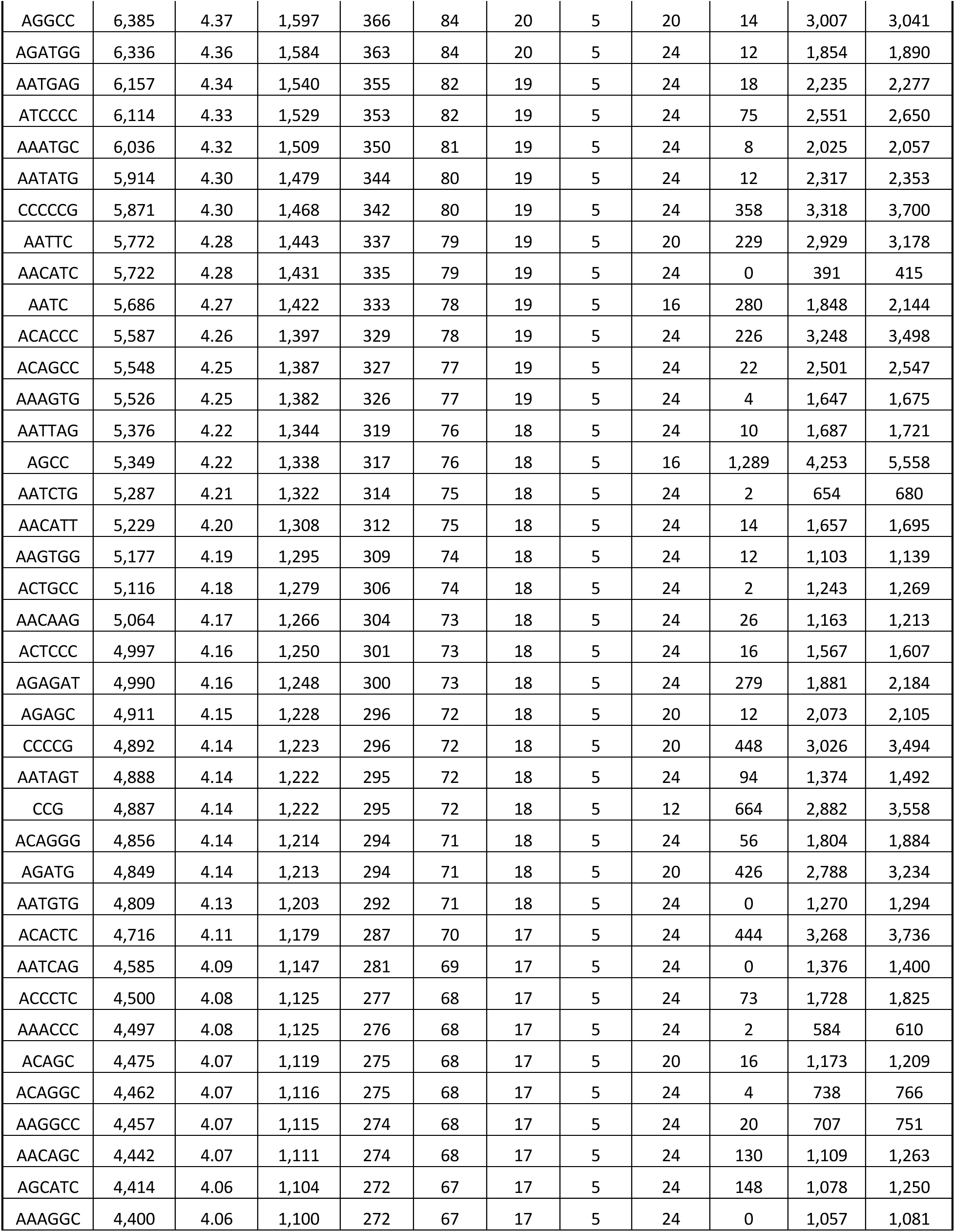
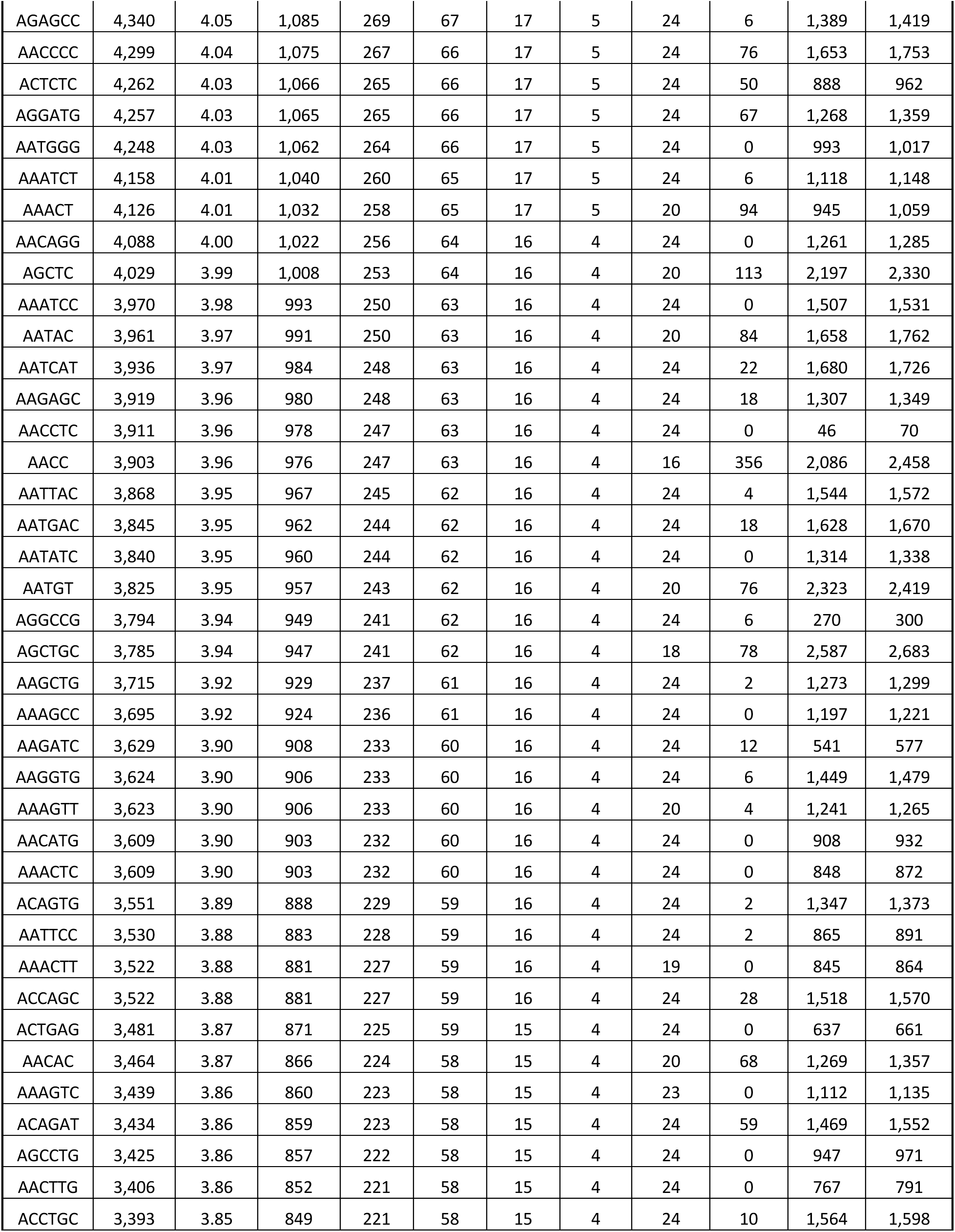
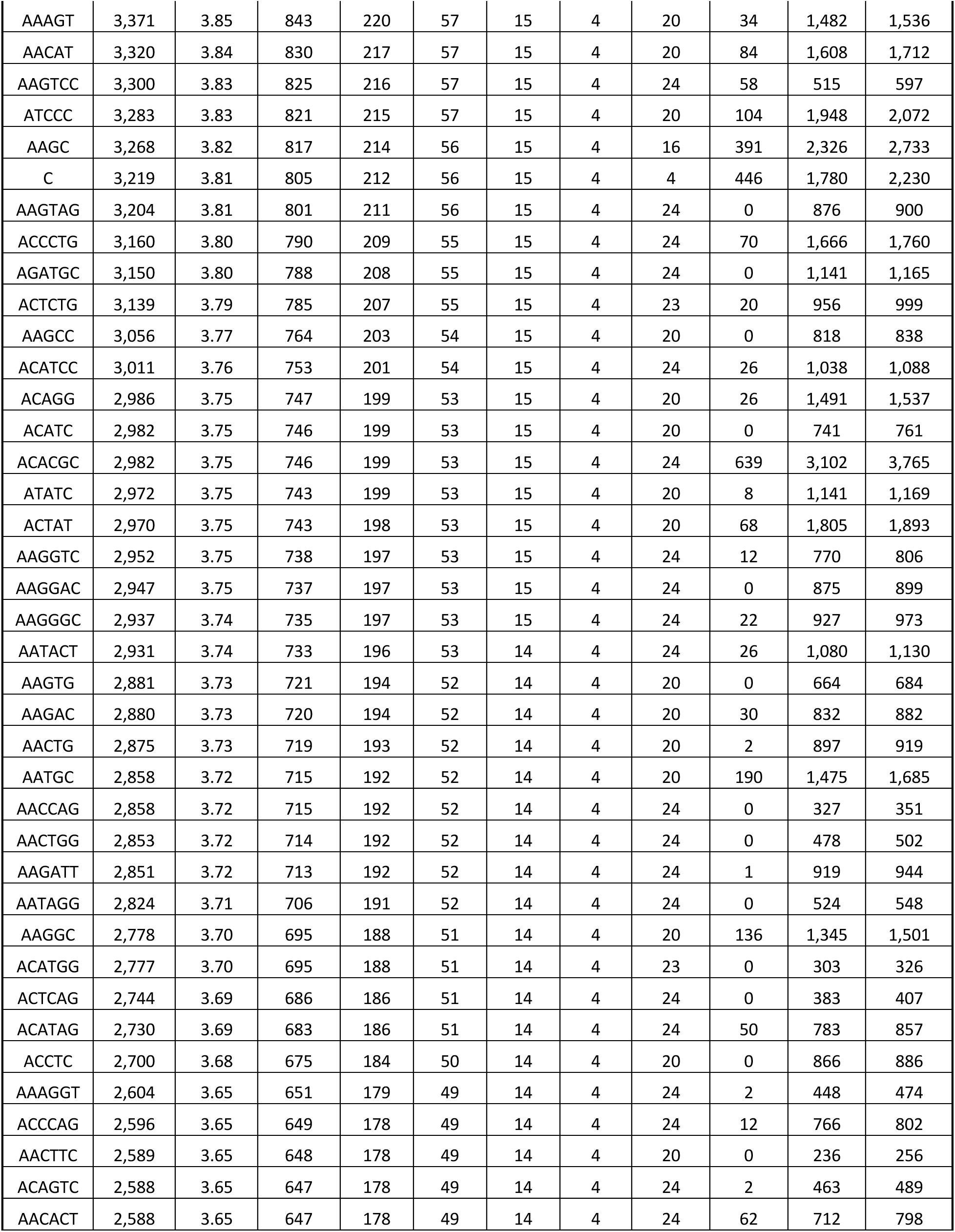
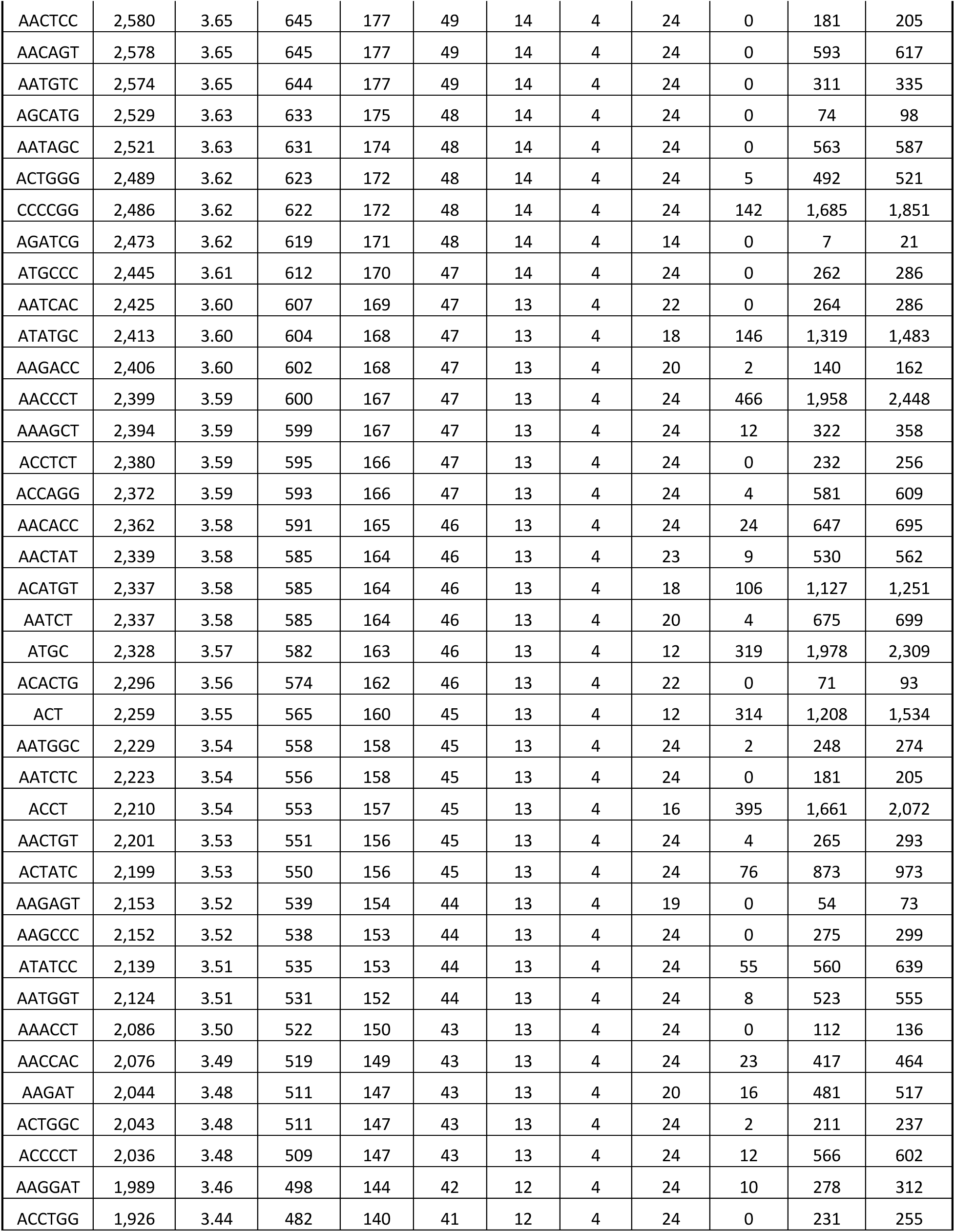
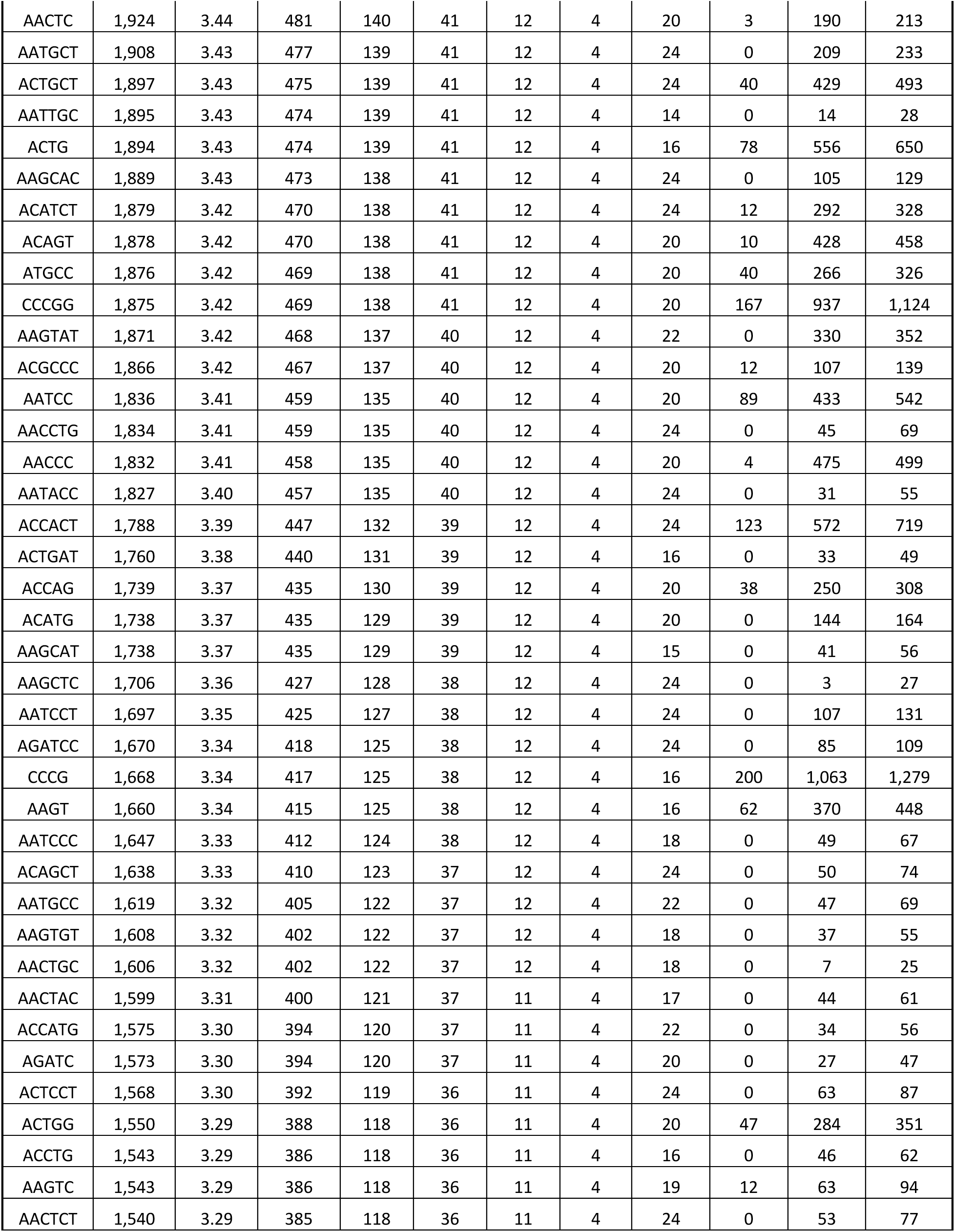
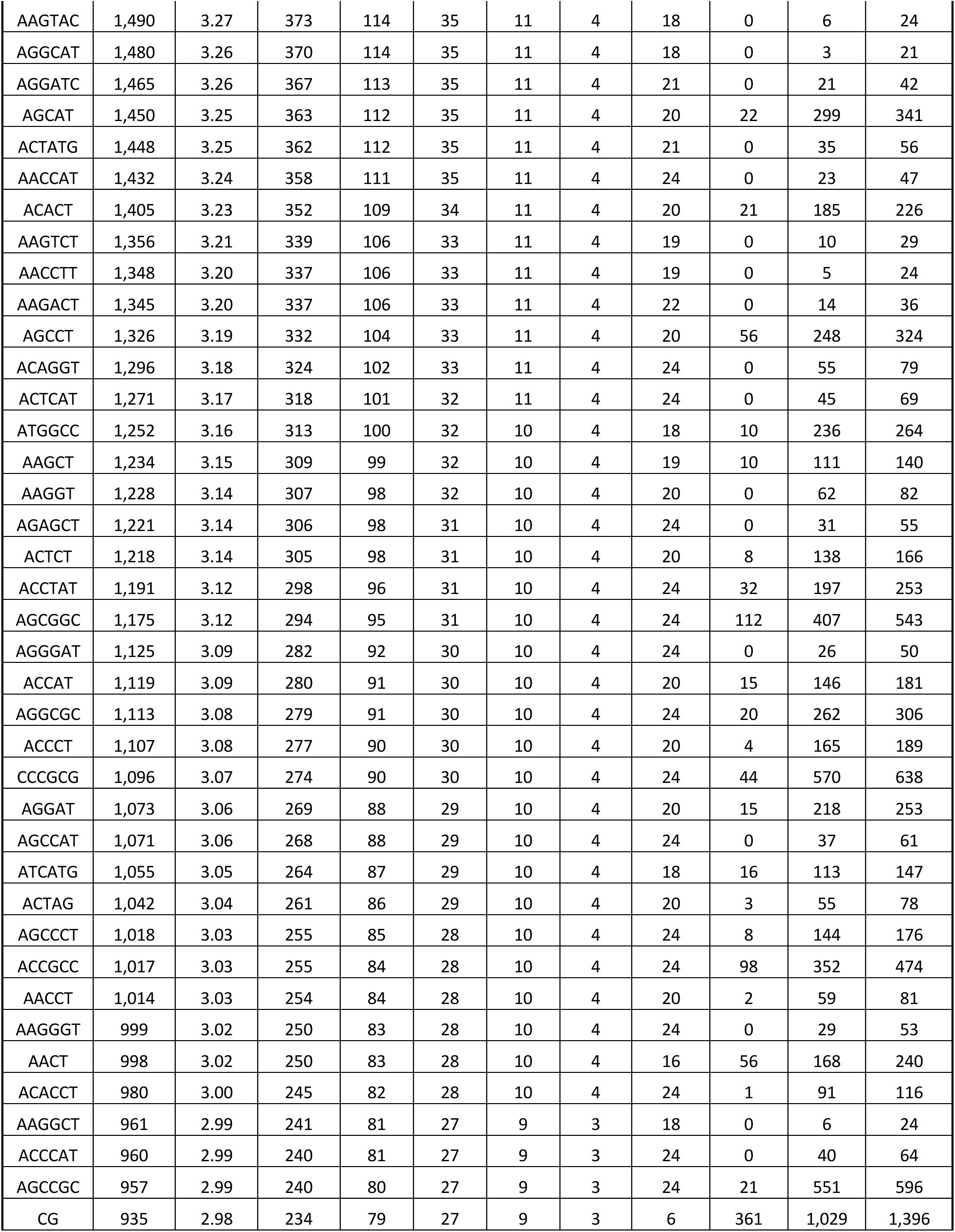
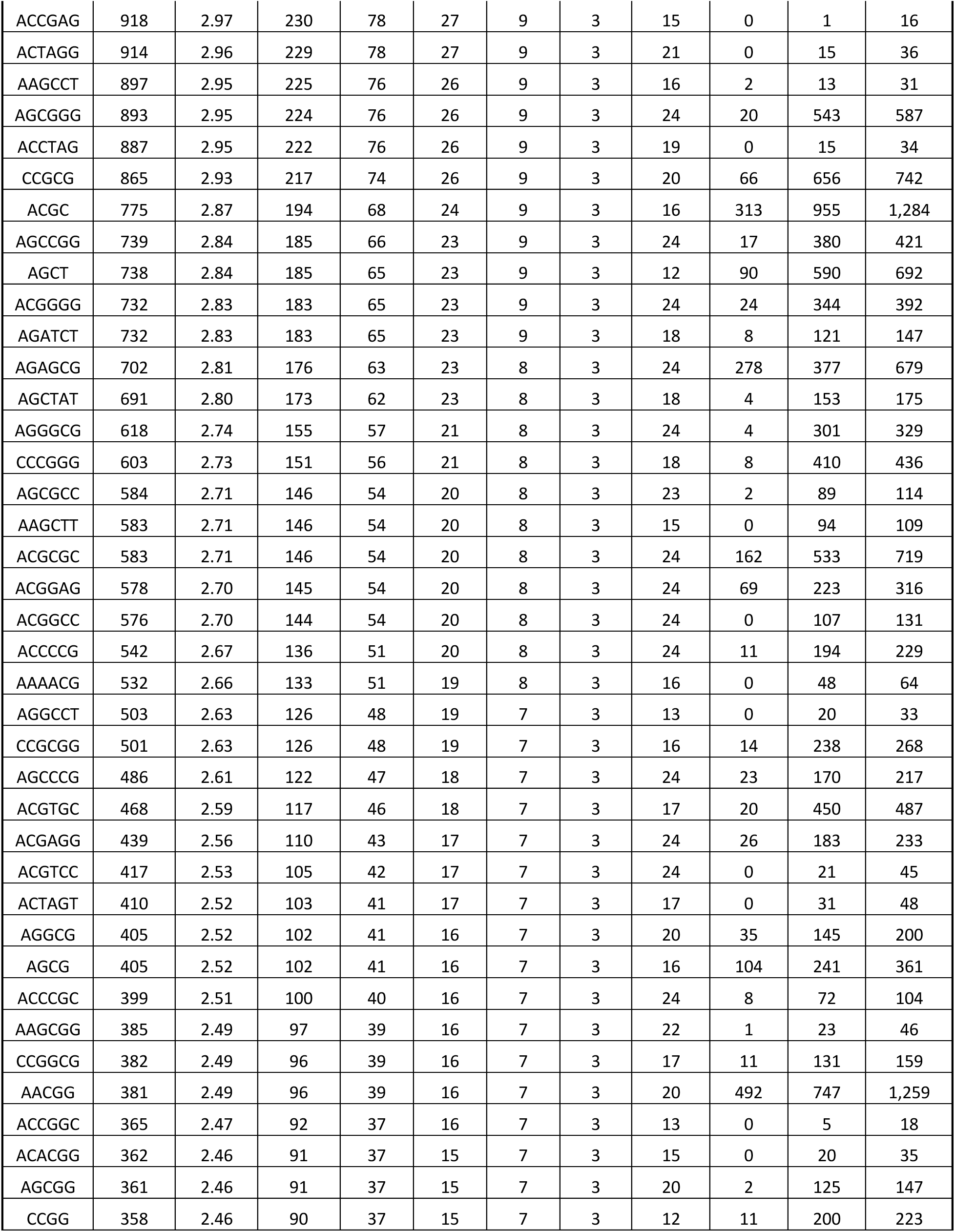
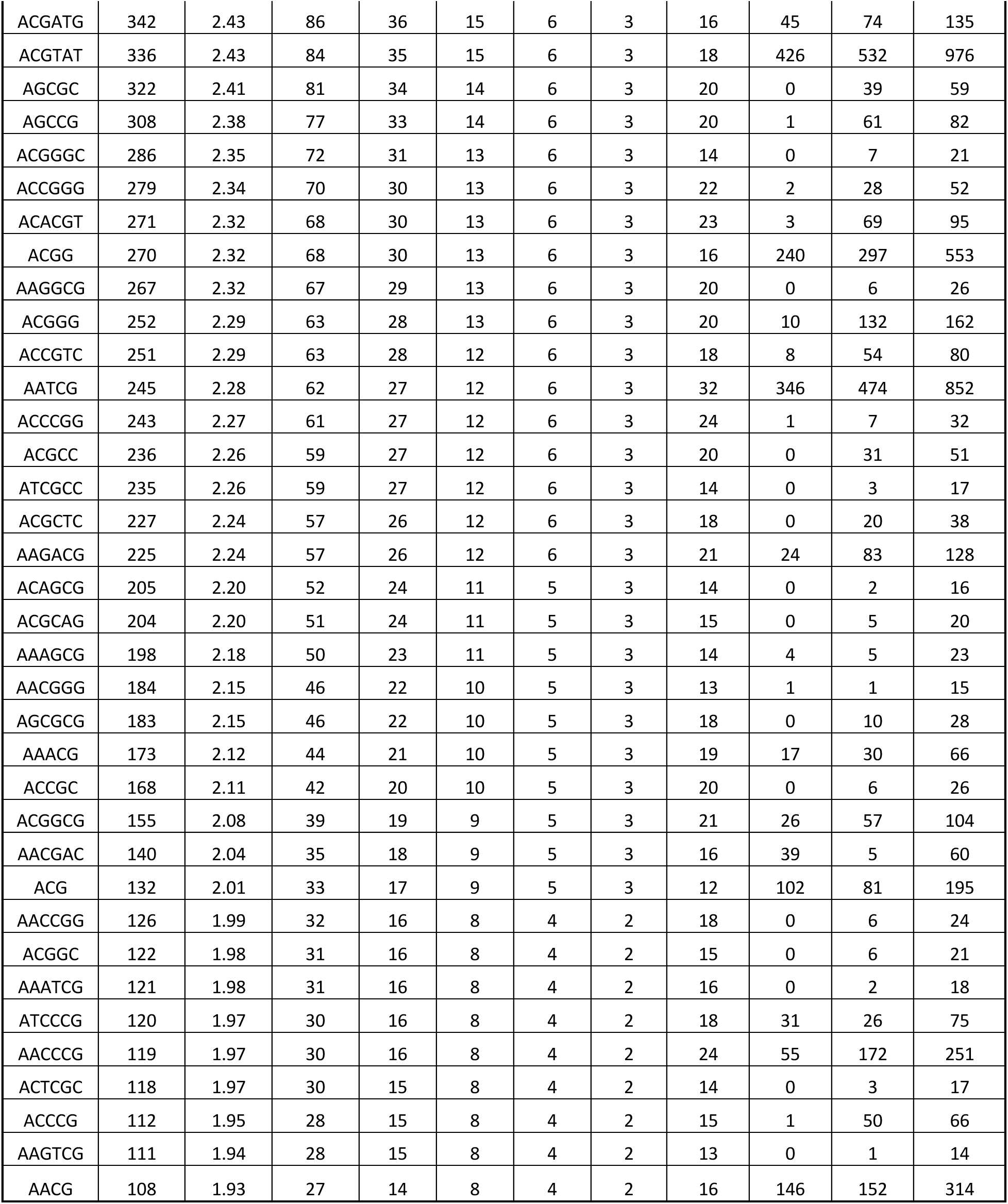
SSR-clouds construction summary.

**Table S2.**
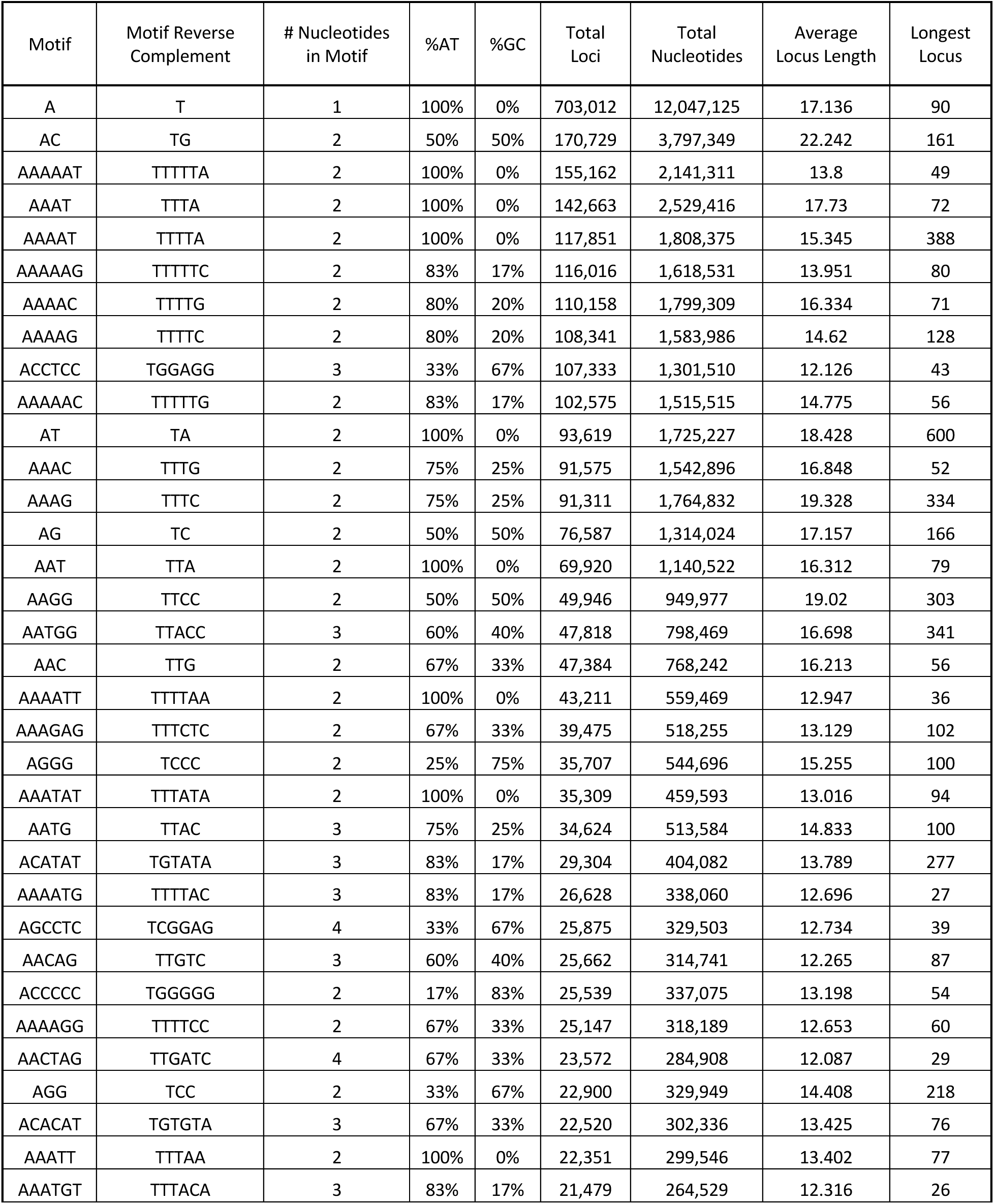
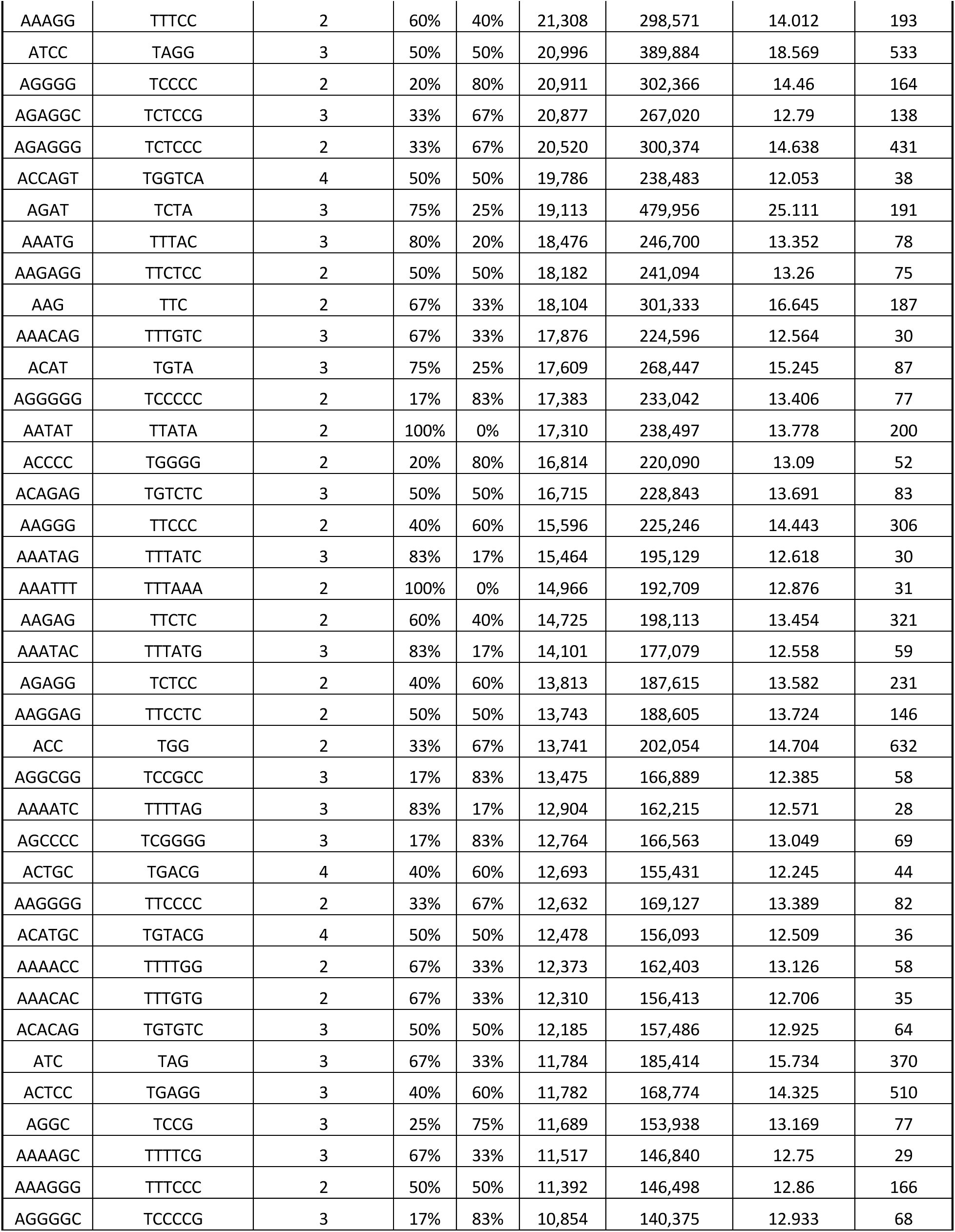
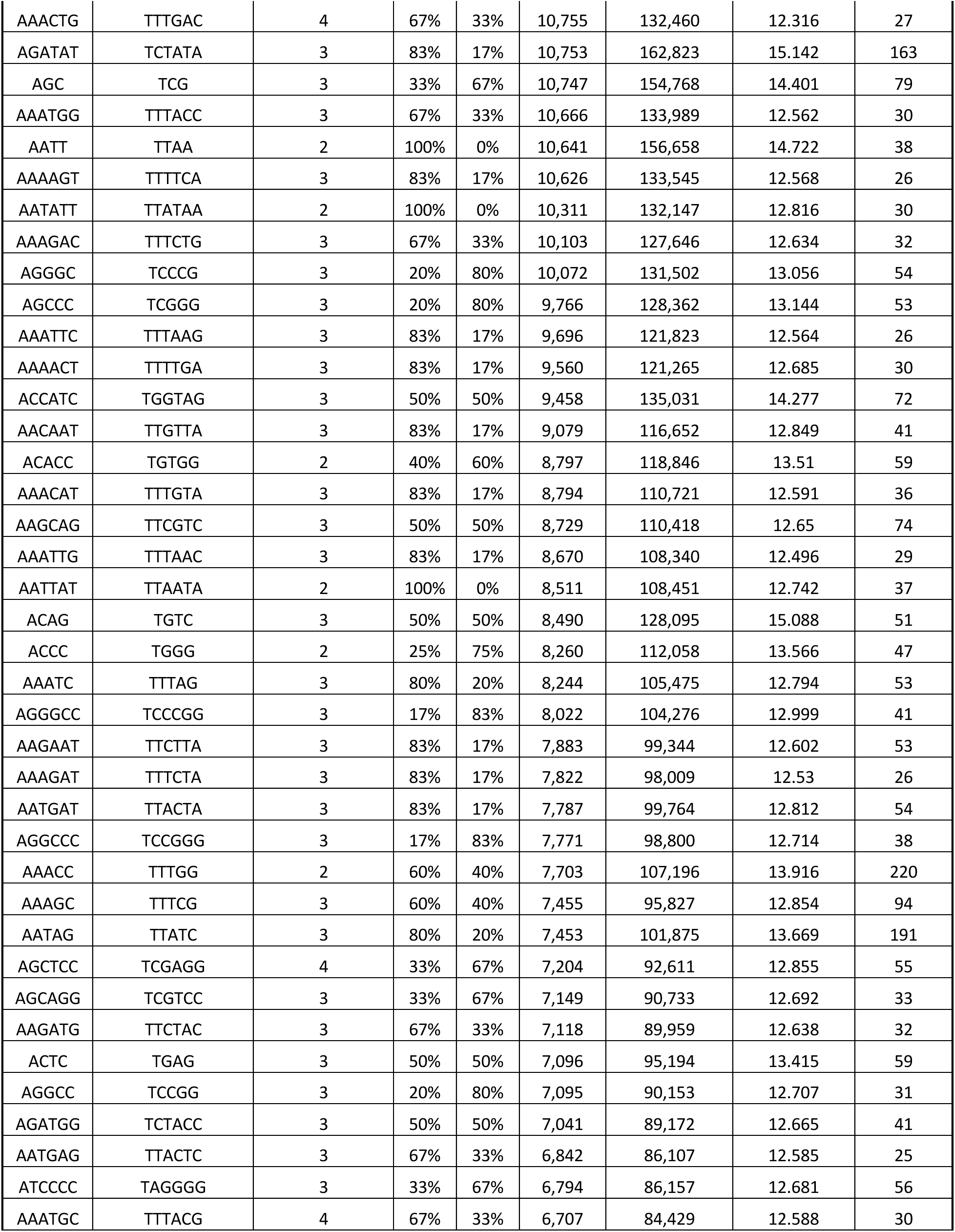
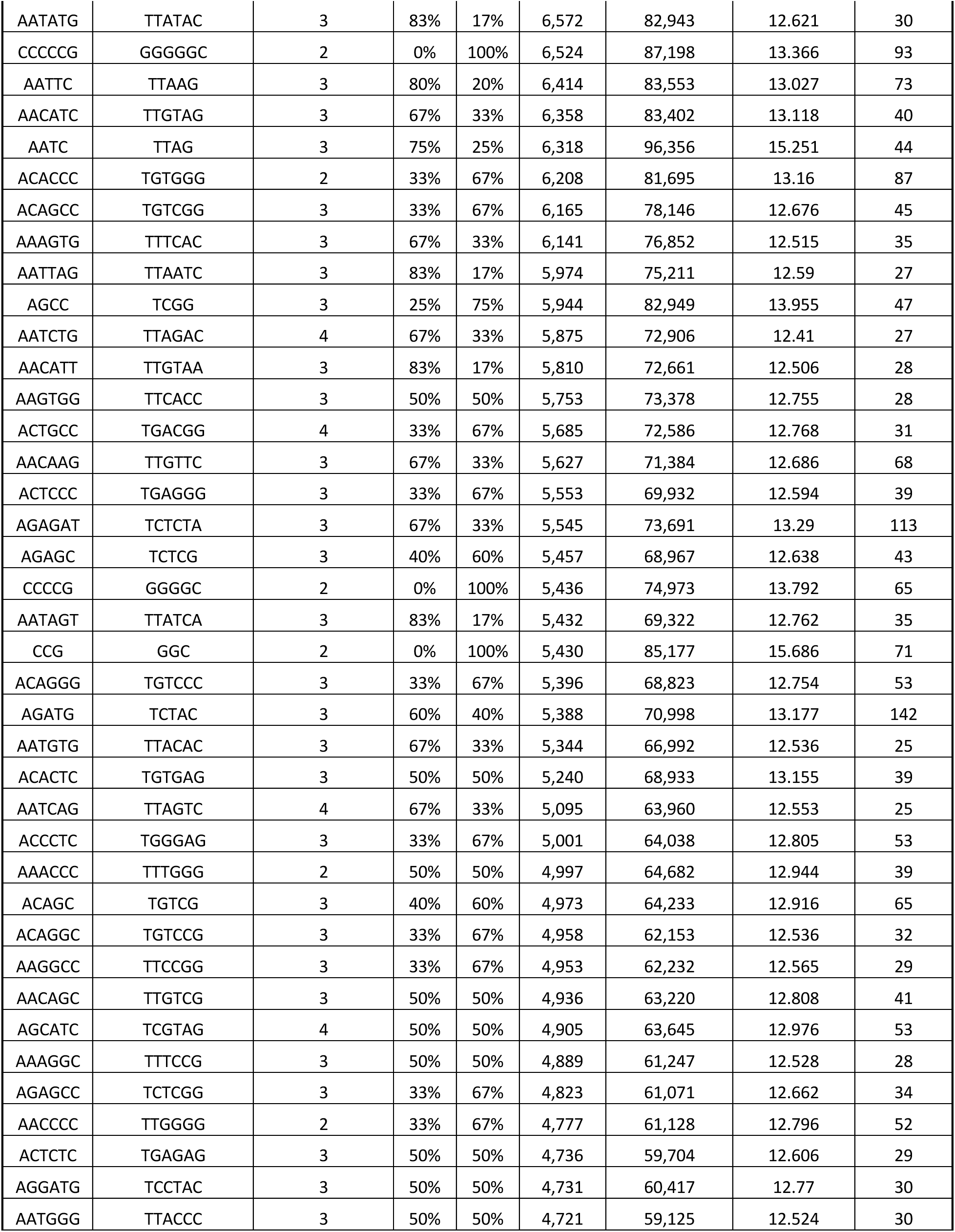
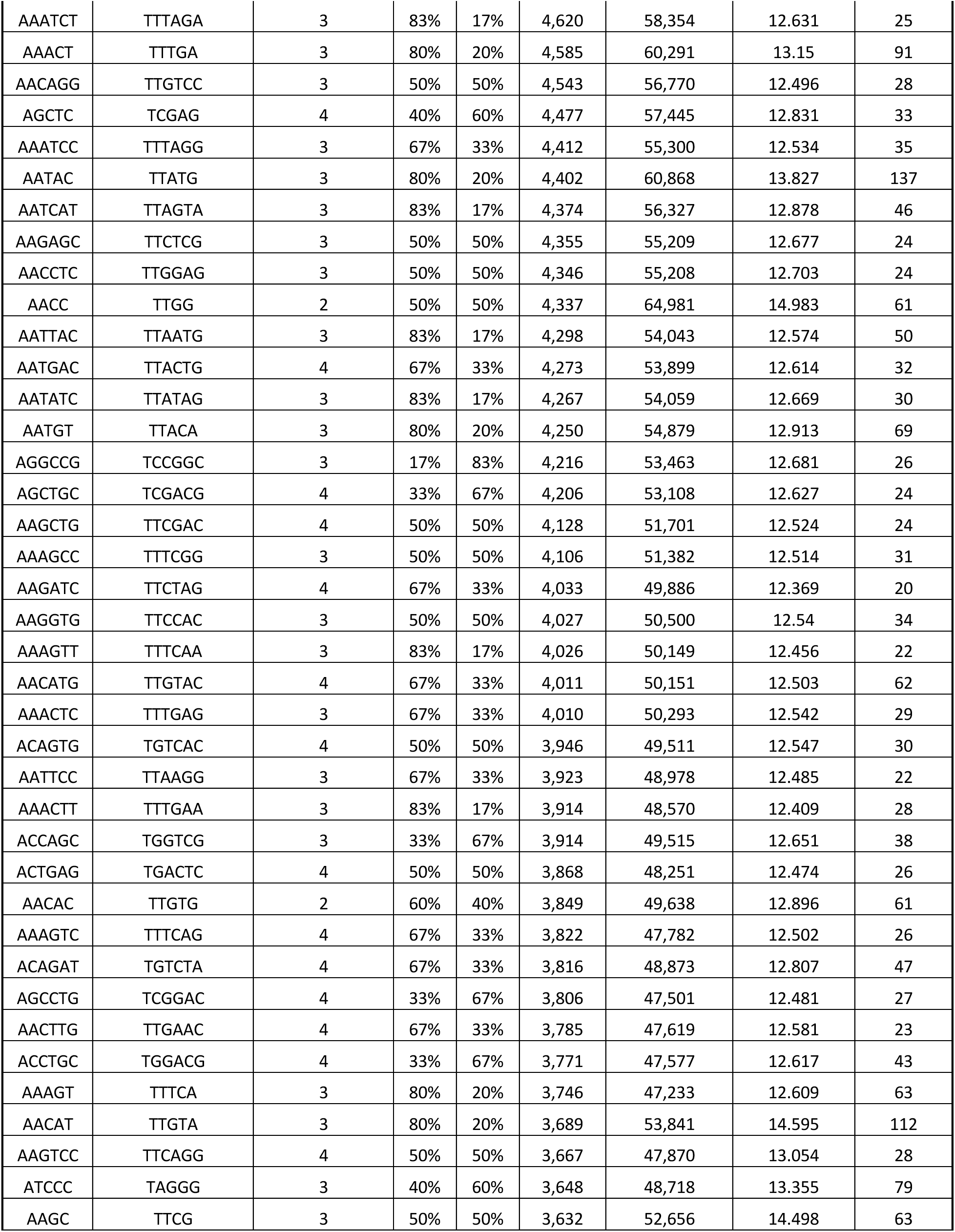
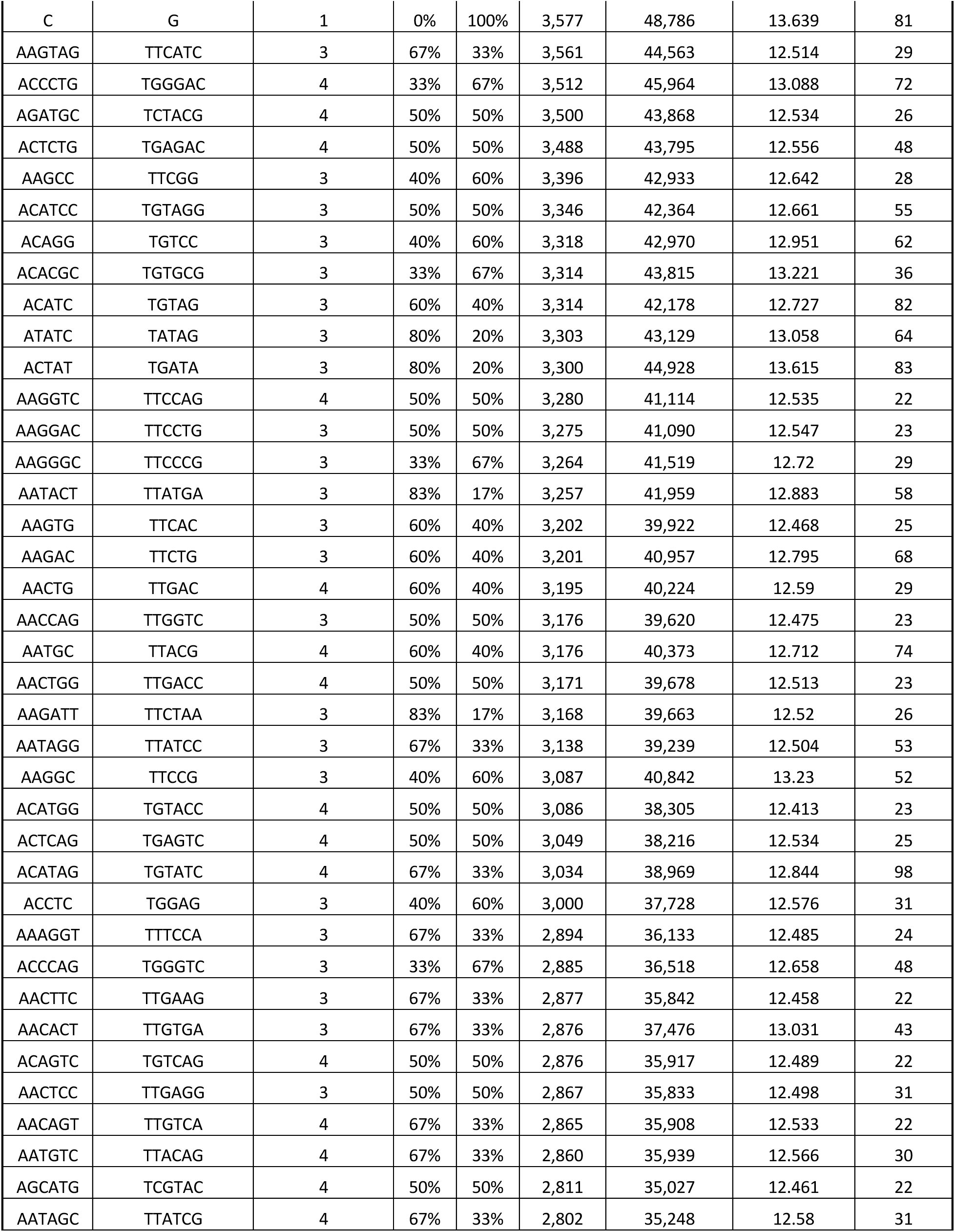
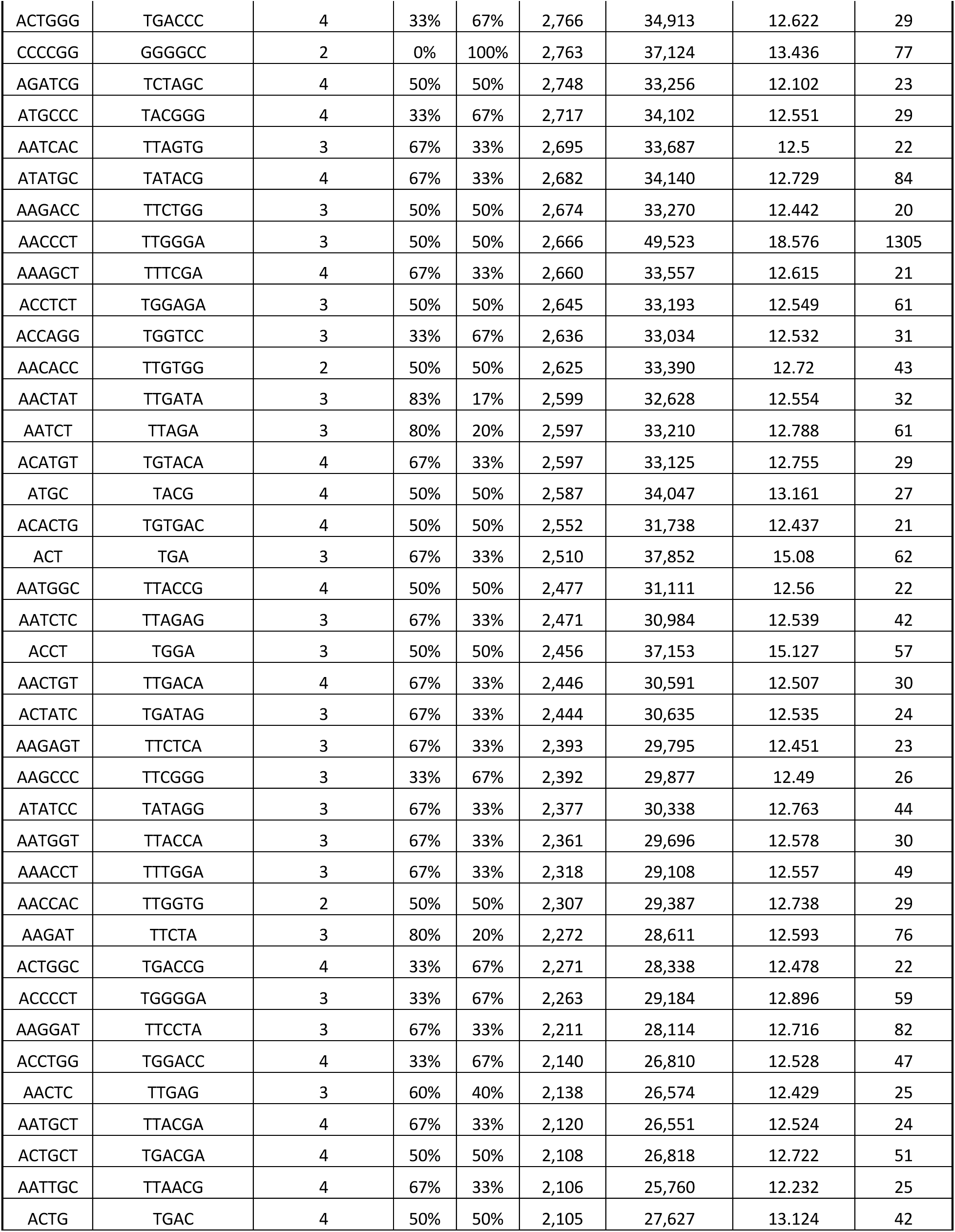
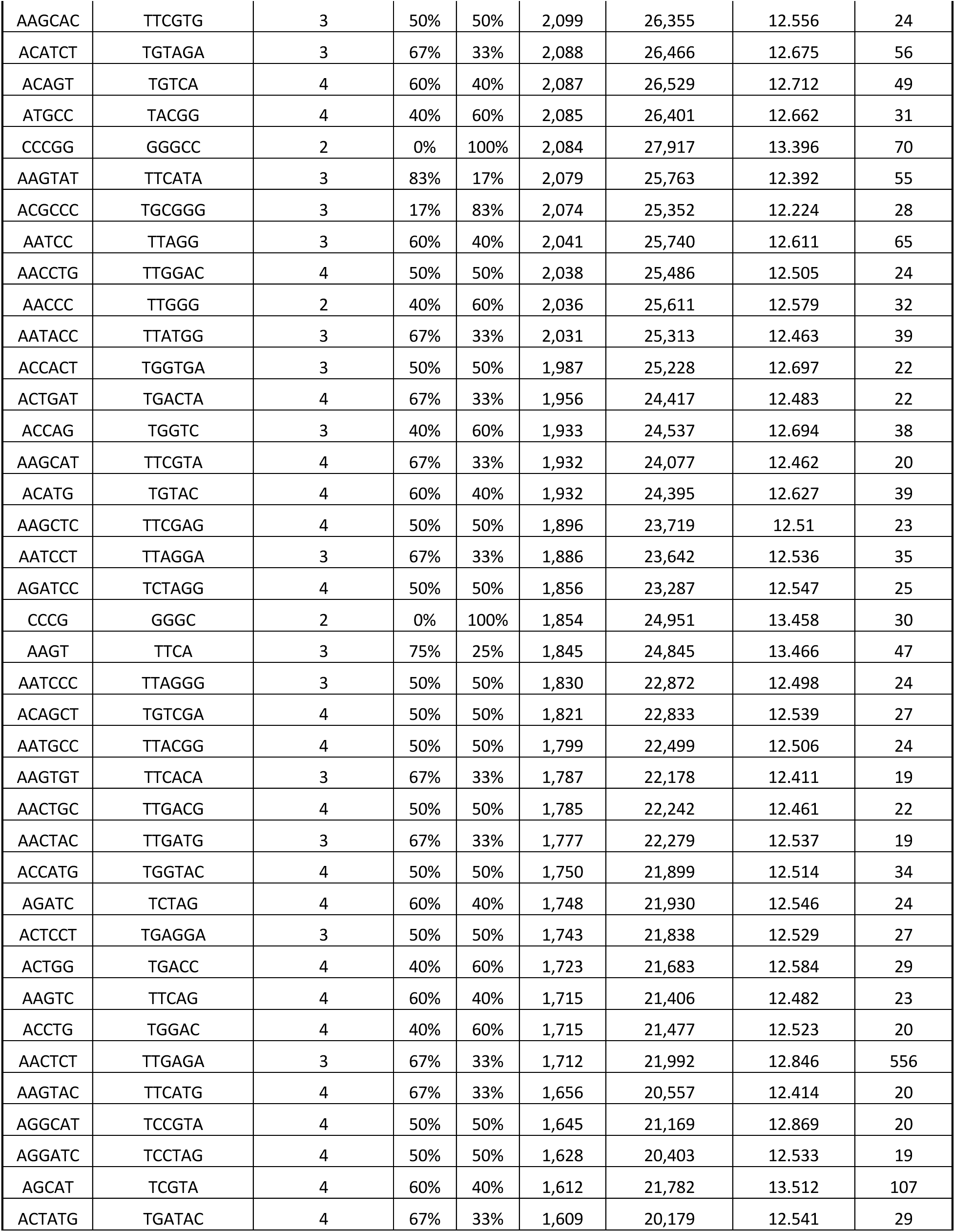
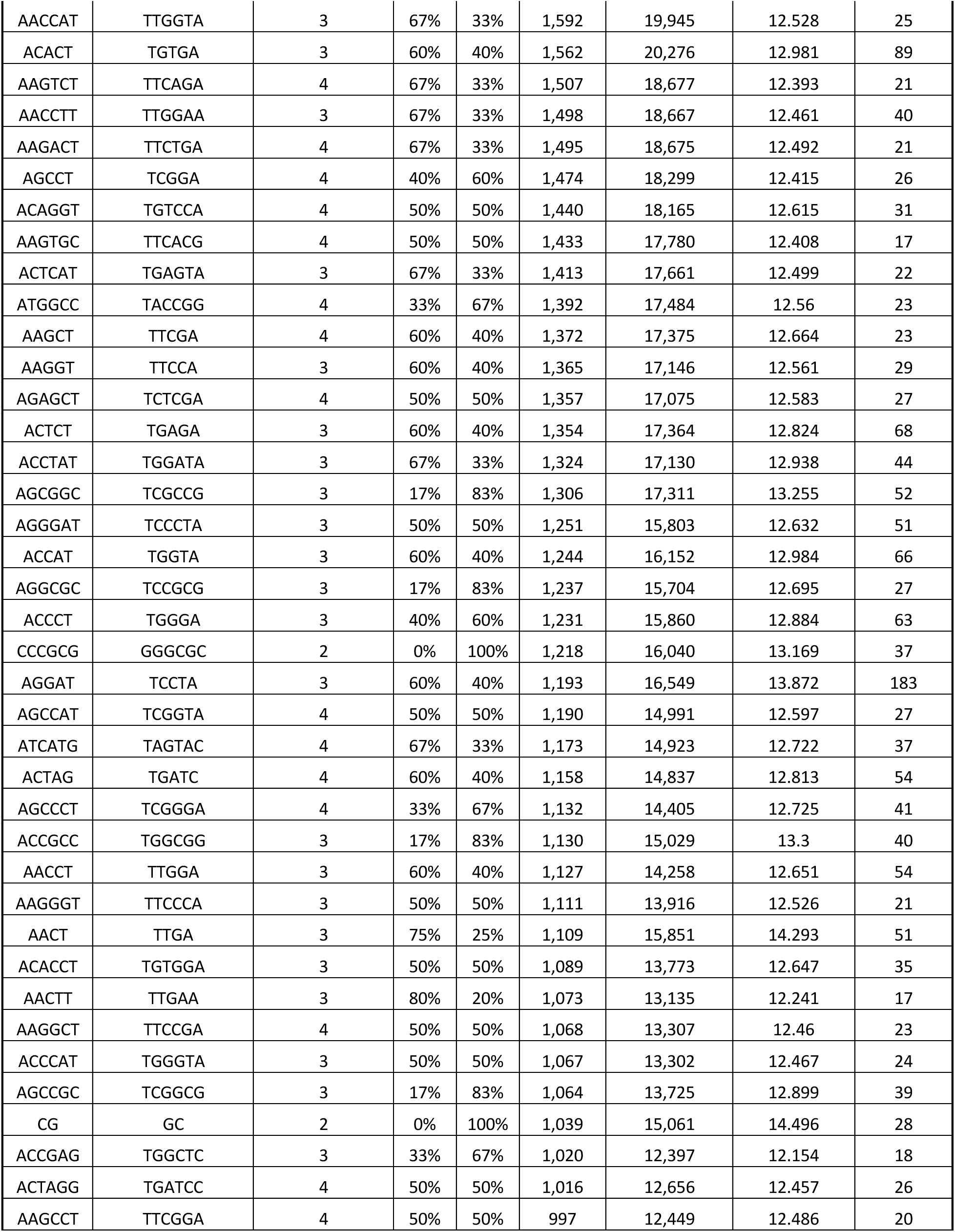
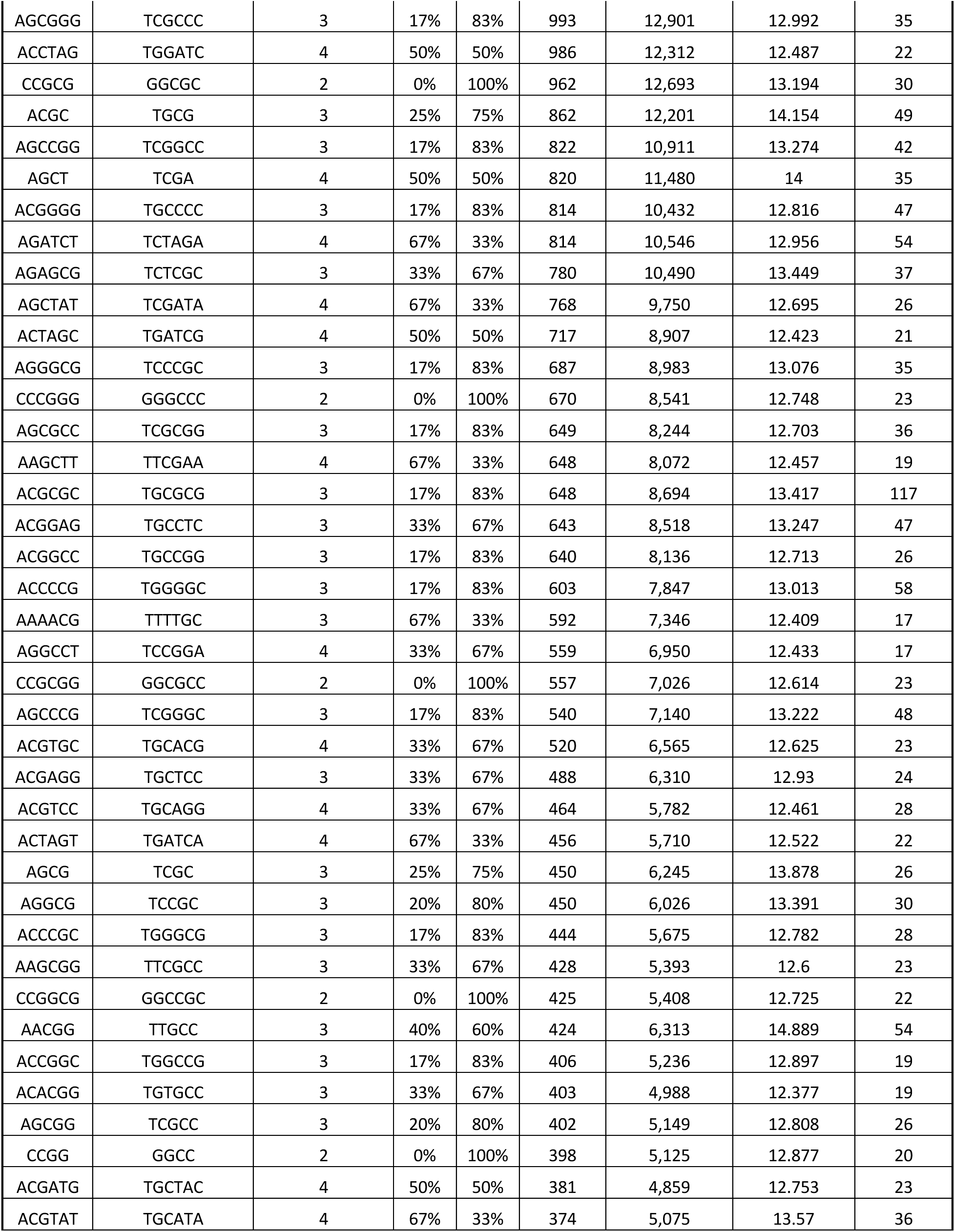
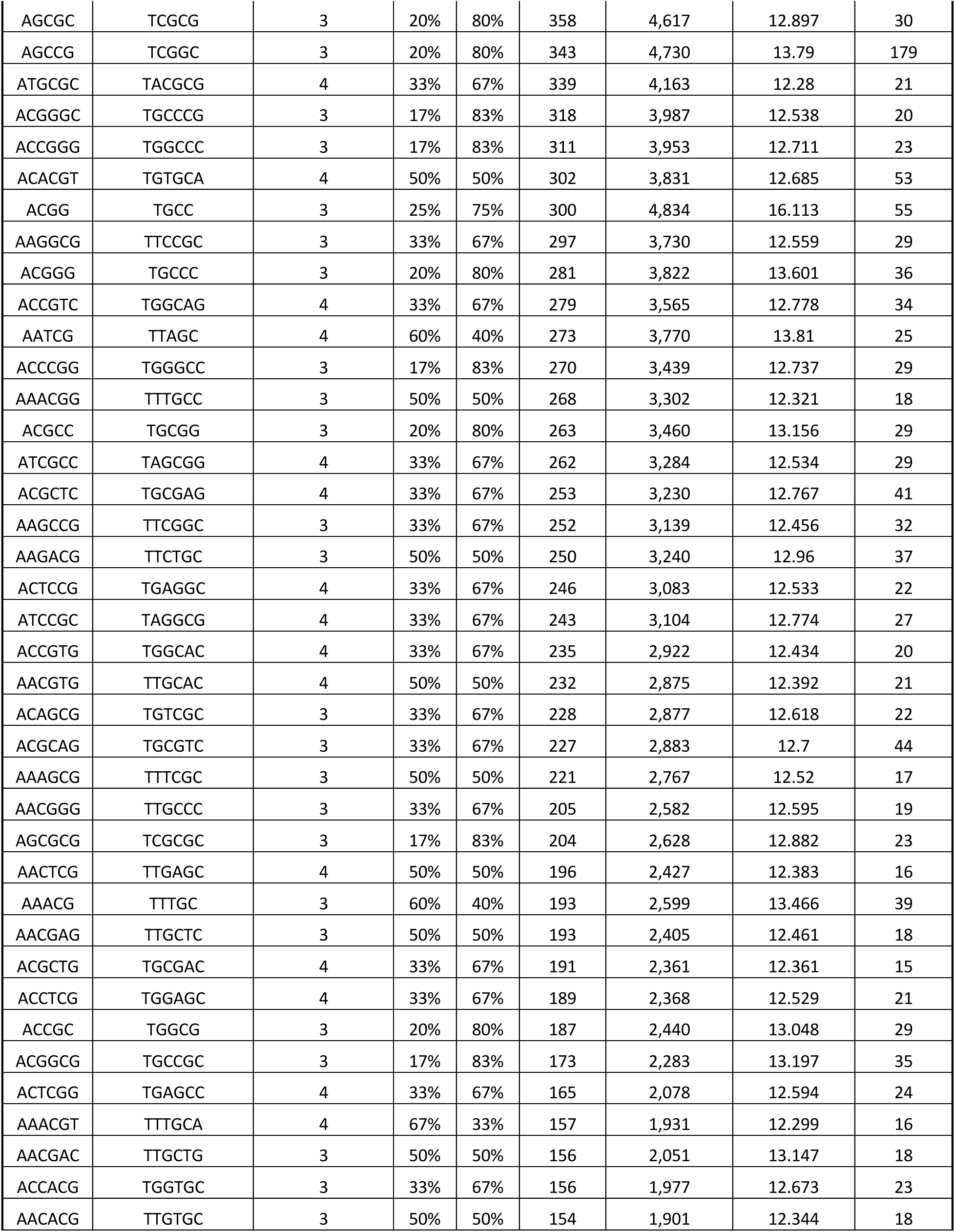
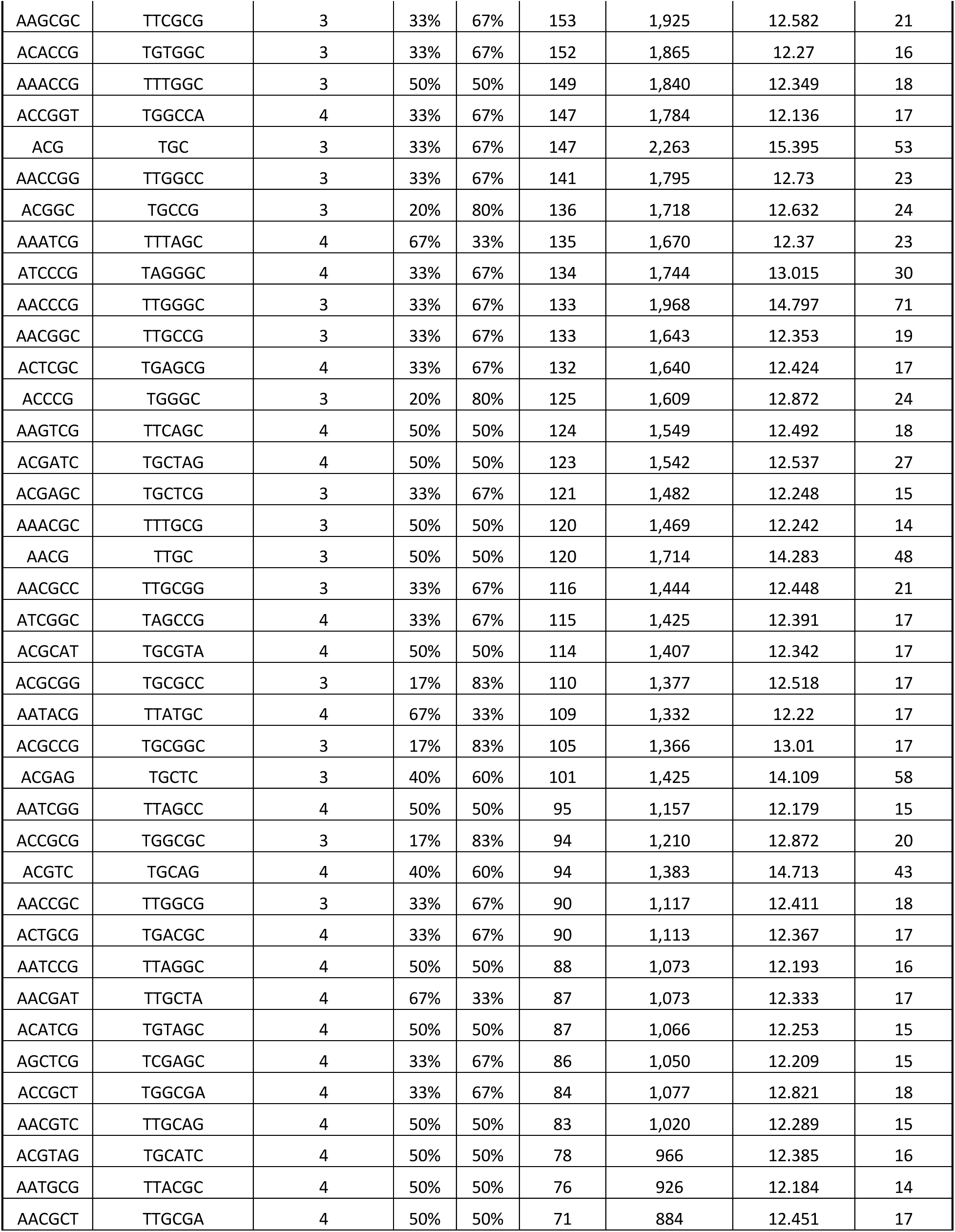
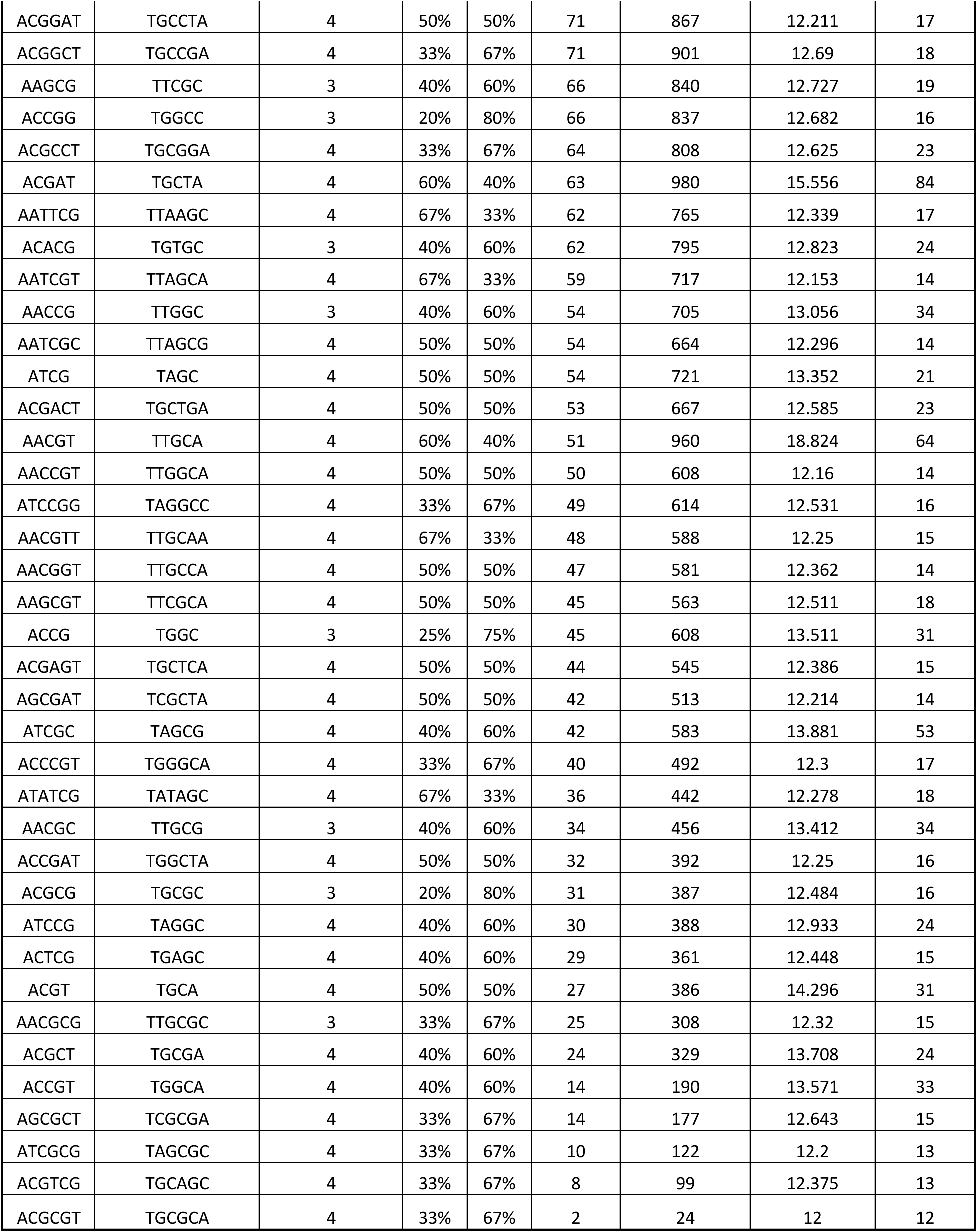
Summary statistics of perfect SSR loci in hg38 for each SSR family.

